# Host cell traversal by *Plasmodium* parasites is essential for sterilizing hepatic cellular immunity

**DOI:** 10.64898/2026.03.20.713177

**Authors:** Ana Rodrigues, António M. Mendes, Rute Gonçalves, Helena Nunes-Cabaço, Sofia Marques, Nuno Valente-Leal, Cristina Ferreira, Marc Veldhoen, Miguel Prudêncio, Maria M. Mota, Ângelo Ferreira Chora

## Abstract

An effective vaccine capable of inducing sterile protection against *Plasmodium*, the causative agent of malaria, is critical to aid global eradication. Whole-organism vaccines using liver-infective sporozoites provide high levels of sterile protection against pre-erythrocytic infection. Yet, determinants of sporozoite immunogenicity remain poorly characterized. Using rodent models of vaccination, we demonstrate that the ability of *Plasmodium* sporozoites to actively migrate through multiple host cells prior to infecting hepatocytes is required for sterilizing immunity, regardless of the time of intrahepatic development of immunizing parasites. We further establish that host cell traversal is sufficient to trigger robust protection against *Plasmodium* hepatic infection. Impaired cell traversal precludes protective liver-resident memory CD8 T cell responses following vaccination, but not the production of anti-plasmodial antibodies. Our findings challenge the prevailing notion that intrahepatic parasite development is the sole determinant of whole-sporozoite vaccination-induced protection, and highlight parasite behavior traits as critical immunogenic events shaping sterilizing cellular immunity against *Plasmodium* liver stages.

## INTRODUCTION

Despite major progress in malaria control over the past two decades, *Plasmodium* infection-associated disease remains a leading cause of morbidity and mortality due to communicable diseases worldwide. Nearly half of the globe population lives at risk of *Plasmodium* infection, with the World Health Organization reporting over 263 million cases and 597,000 deaths in 2023 alone^1^. In malaria endemic regions, natural exposure to *Plasmodium* is non-sterilizing^2^. This persistent susceptibility to re-infection highlights the immunological challenges faced when developing a malaria vaccine capable of conferring complete and durable protection.

From all malaria vaccination strategies targeting the different stages of parasite development, those directed against the pre-erythrocytic stages of *Plasmodium* infection are especially promising^3^. During this obligatory phase of infection, which precedes the onset of blood stage disease, sporozoites (spz) are inoculated by an infected anopheline mosquito, enter the bloodstream, and travel to the liver. There, spz traverse multiple non-parenchymal and parenchymal cells by repetitively breaching cell membranes, migrating through the cytoplasm, and exiting without establishing infection, in a process termed cell traversal (CT)^4,5^.

To date, two malaria subunit vaccines directed against pre-erythrocytic antigens have been licensed for use in endemic settings. While these vaccines reduce disease burden, they do not consistently prevent infection^6–9^, a key requirement for eradication. In contrast, whole-sporozoite (W-SPZ) vaccination strategies employing radiation- or genetically-attenuated spz, or fully infectious spz under chemoprophylaxis induce sterilizing immunity in both humans^10,11^ and rodent models^12^. The current view is that this protection largely depends on the capacity of immunizing parasites to infect hepatocytes and sustain prolonged, or even complete, intrahepatic development^13–15^. However, sterile protection is also achieved with attenuated parasites whose development arrests early after infection^10,16,17^, suggesting that events preceding hepatocyte infection may contribute to vaccine efficacy.

Here, we show that host cell traversal by immunizing spz during the initial steps of infection plays a critical role in shaping protective immunity. We demonstrate that CT is not only required but also sufficient to induce sterilizing protection following W-SPZ vaccination in rodents by promoting the establishment of liver-resident memory CD8 T cells. Our findings open new avenues for rational vaccine design and highlight the need to consider parasite behavior traits, and not only antigenic load and diversity, for optimizing protective immunity against *Plasmodium*.

## RESULTS

### Cell traversal is required for sterile protection following W-SPZ vaccination

We sought to compare the immunogenicity of CT-competent *P. berghei* (*Pb*) wild-type (*Pb*WT) parasites to that of CT-deficient parasites lacking the *sporozoite microneme protein essential for cell traversal 1* gene (*PbSpect1*^-^)^18^. To control for potential immunogenic effects of the SPECT1 protein itself, we generated a distinct CT motility-deficient parasite line lacking the *perforin-like protein 1* gene (*PbPlp1*^-^) (**Extended Data Fig. 1a,b**). As previously reported^19^, *PbPlp1*^-^ parasites failed to traverse HepG2 cells in *in vitro* cell wounding assays (**Extended Data Fig. 1c**), while displaying comparable intrahepatic development to their wild-type counterparts (**Extended Data Fig. 1d**). Both CT-deficient parasite lines exhibit reduced infectivity *in vivo*^18,19^, complicating the direct comparison of their protective efficacy relative to *Pb*WT parasites at equivalent liver infection loads. To overcome this, the liver loads of increasing inocula of *PbSpect1*^-^ and *PbPlp1*^-^spz were compared to that resulting from the inoculation of 10^4^ *Pb*WT spz, revealing that equivalent liver loads at 16 hours post infection (h.p.i.) required a 15:1 inoculation ratio of *PbSpect1*^-^ or *PbPlp1*^-^ radiation-attenuated sporozoites (RAS) relative to *Pb*WT RAS (**Fig. 1a,b**). Using this standardized dose, we employed a well-established prime–boost–boost RAS immunization protocol in C57BL/6 mice^20^ (**Fig. 1c**) to evaluate the protective efficacy of each *Pb* line in the 12 day-period following challenge with fully infectious, luciferase-expressing *Pb*WT parasites. *In vivo* bioluminescence imaging at 48 hours post challenge (h.p.c.) revealed that non-immunized (NI) mice harbored metabolically active parasites in the liver, whereas *Pb*WT RAS-immunized mice had no detectable bioluminescence activity (**Fig. 1d** and **Extended Data Fig. 1e**). Contrastingly, both *PbSpect1*^-^ and *PbPlp1^-^* RAS-immunized mice displayed higher bioluminescence activity than *Pb*WT RAS-immunized mice, albeit significantly reduced compared to NI mice (**Fig. 1d** and **Extended Data Fig. 1e**). Sterile protection, assessed by the absence of blood stage parasites following challenge by bioluminescence, was observed in 82.50±4.53 of the *Pb*WT RAS-immunized cohort (**Fig. 1e**), whereas blood infection became patent in 100% of *PbSpect1*^-^ and 86.67±13.33% of *PbPlp1^-^* RAS-immunized mice on days 4.70±0.18 and 4.23±1.3 after challenge, respectively (**Fig. 1e**). Erythrocytic infection became patent in unprotected *Pb*WT RAS-immunized mice on day 6.14±0.63 (**Fig. 1e**). The decreased number of parasites seeding the blood stage of infection led to the delayed progression of parasitemia in unprotected mice from all immunized cohorts (**Extended Data Fig. 1f**) with varying levels of protection from experimental cerebral malaria (ECM) (**Extended Data Fig. 1g**). Several studies suggest that the timing of intrahepatic parasite arrest plays a major role in the potency of the immune response elicited by vaccination, with longer development of immunizing parasites potentially exposing the host to a wider breadth and quantity of antigens^21,22^. To assess if CT remained relevant for sterile protection following immunization regimens allowing for extended or even complete intrahepatic development of immunizing spz, we employed two immunization regimens using infectious spz under azithromycin or chloroquine chemoprophylaxis, i.e., CPS-AZ^23^ and CPS-CQ^24^ (**Fig. 1f**). We validated that the previously established 15:1 ratio of *PbSpect1*^-^ to *Pb*WT spz led to comparable parasite loads at late stages of hepatic infection (**Fig. 1g**), with both lines developing to a similar extent (**Fig. 1h**). *Pb*WT-immunized mice under both CPS immunization regimens had no detectable metabolically active parasites in the liver at 48 h.p.c. (**Fig. 1i** and **Extended Data Fig. 1h,k**), with 100% and 80±11.55% of immunized mice remaining blood stage parasite-free for the 12 day-period following challenge with *Pb*WT spz (**Fig. 1j,k**). In contrast, *PbSpect1*^-^ CPS-AZ- and CPS-CQ-immunized cohorts exhibited only partial reductions in liver stage parasite load (**Fig. 1i** and **Extended Data Fig. 1h,k**) and failed to achieve sterile protection, with blood stage infection becoming patent on days 5.10±0.18 and 4.0±0.15 after challenge, respectively (**Fig. 1j,k**). Unprotected mice, irrespective of receiving CT-competent or CT-deficient immunizing parasites, exhibited delayed parasitemia (**Extended Data Fig. 1i,l**) and were protected from ECM (**Extended Data Fig. 1j,m**). Notably, azithromycin also prevented ECM in NI controls (**Extended Data Fig. 1j**), suggesting that it can counter severe neurological manifestations of malaria. Collectively, these data demonstrate that host cell traversal by immunizing sporozoites is a critical determinant of pre-erythrocytic sterilizing immunity following W-SPZ vaccination, regardless of the duration of intrahepatic parasite development.

**Fig. 1.**
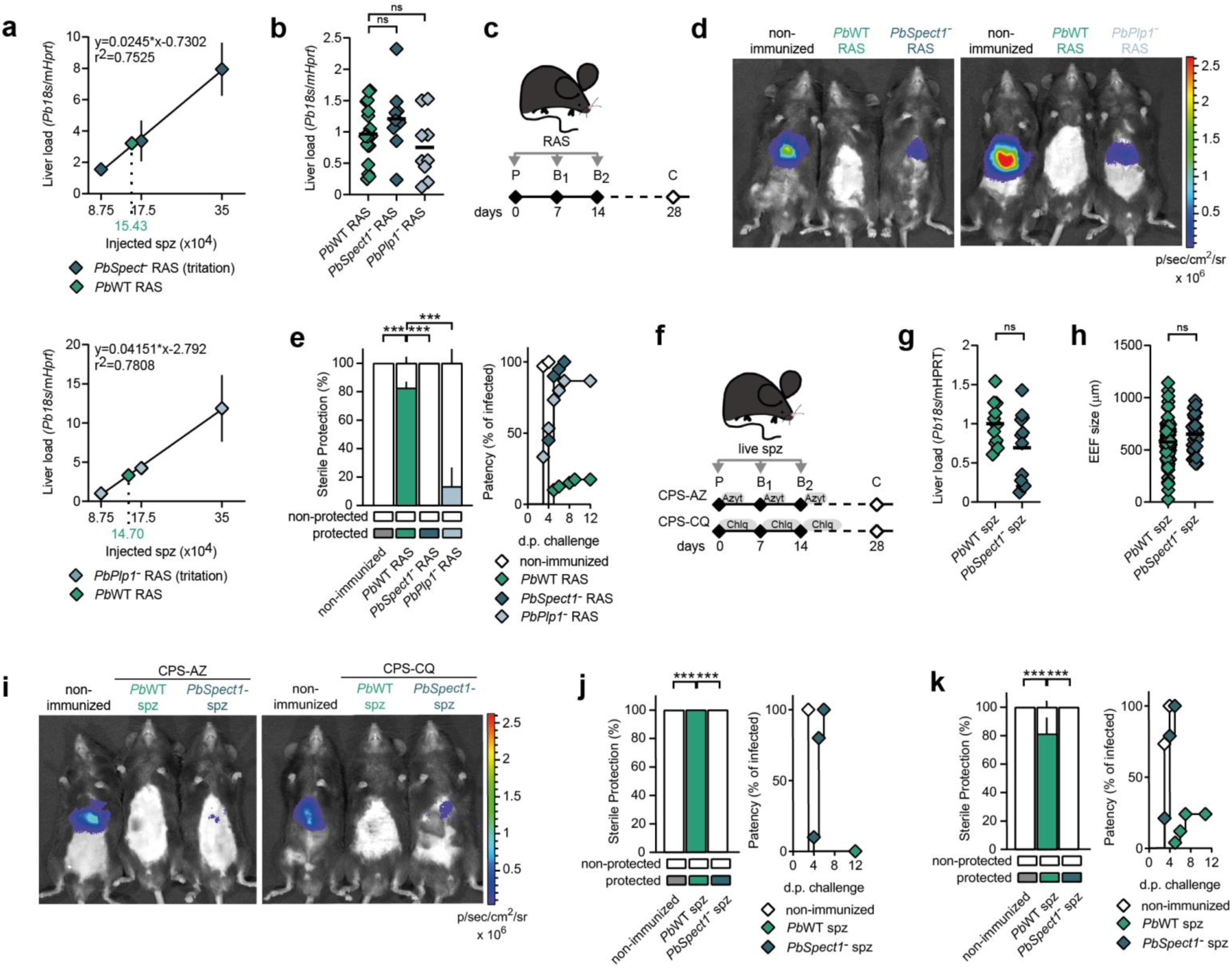
Cell traversal motility of immunizing parasites is required for sterilizing immunity against *Plasmodium* hepatic infection. **a**, Linear regression of *Plasmodium 18s* rRNA levels in the livers of C57BL/6 mice, 16 hours after receiving the indicated amounts of (upper panel) *PbSpect1^-^* (n=3 per dose) or (bottom panel) *PbPlp1^-^* (n=3 per dose) or 10^4^ *Pb*WT (n=3) spz. Dotted lines indicate the estimated number of *PbSpect1^-^* or *PbPlp1^-^* spz required to achieve liver infection loads equivalent to that yielded by injection of 10^4^ *Pb*WT. **b**, Quantification of *Plasmodium 18s* rRNA levels in the livers of C57BL/6 mice, 16 hours after receiving 10^4^ *Pb*WT (n=20), 15x10^4^ *PbSpect1^-^* (n=10) or 15x10^4^ *PbPlp1^-^* (n=9) RAS (2-4 independent experiments). **c**, Protocol of RAS immunizations and challenge. **d**, Representative luciferase bioluminescence images of non-immunized, and *Pb*WT, *PbSpect1^-^* (left) or *PbPlp1^-^* (right) RAS-immunized C57BL/6 mice, 48 hours after challenge with *Pb*WT spz. **e**, Sterile protection, represented as percentage of blood stage parasite-free mice (left), and time to patency (right) of non-immunized (n=35), or *Pb*WT (n=40), *PbSpect1^-^* (n=20) or *PbPlp1^-^*(n=15) RAS-immunized C57BL/6 mice in the 12-days period following challenge with *Pb*WT spz (3-8 independent experiments). **f**, Protocol of immunizations with live spz under chemoprophylaxis with azithromycin (CPS-AZ) or chloroquine (CPS-CQ) and challenge. **g**, **h** Quantification of (**g**) *Plasmodium 18s* rRNA levels and (**h**) exo-erythrocytic forms (EEF) size in the livers of C57BL/6 mice, 42 hours after inoculation of 10^4^ *Pb*WT (n=11) or 15x10^4^ *PbSpect1^-^*(n=12) live spz (3 independent experiments). **i**, Representative luciferase bioluminescence image of non-immunized, and *Pb*WT or *PbSpect1^-^* CPS-AZ- or CPS-CQ-immunized C57BL/6 mice, 48 hours after challenge with *Pb*WT spz. **j**, **k** Sterile protection, represented as percentage of blood stage parasite-free mice (left), and time to patency (right) of non-immunized, and *Pb*WT or *PbSpect1^-^* CPS-immunized C57BL/6 mice under (**j**) AZ (n=5, n=15 and n=10, respectively) or (**k**) CQ (n=15, n=25 and n=19, respectively) coverage in the 12-days period following challenge with *Pb*WT spz (1-4 independent experiments). **a**, Data are represented as mean ± SEM, with a simple linear regression connecting all data points. **b**, **g**, and **h**, Data (Mann-Whitney test) are represented as dot plots, black line represents mean. **e**, **j** and **k**; left panels, Data (Mann-Whitney test) are represented as bars ± SEM. **e**, **j** and **k**; right panels, Data (Mann-Whitney test) are represented as the mean day to patency. ns – non-significant; * - p<0.05; ** - p<0.01; *** - p<0.001. For complete statistical analysis, please refer to Source Data Fig. 1.

### Cell traversal by immunizing spz is sufficient to establish sterilizing immunity

Next, we interrogated the relative contributions of CT and hepatocyte infection for protection by employing *Pb* parasites lacking the *P36p* gene (*PbP36p*^-^)^25^, which retained *in vivo* CT activity (**Extended Data Fig. 2a**), but exhibited markedly reduced hepatocyte infectivity (**Extended Data Fig. 2b**). Both *Pb*WT and *PbP36p^-^*RAS-immunized mice (**Fig. 1c**) showed no liver bioluminescence at 48 h.p.c., indicating the successful elimination of hepatic parasites (**Fig. 2a** and **Extended Data Fig. 2c**). Consistently, both immunized cohorts exhibited comparable levels of sterile protection (**Fig. 2b**). Delayed blood stage onset in unprotected *Pb*WT and *PbP36p*^-^ RAS-immunized mice (**Fig. 2b** and **Extended Data Fig. 2d**) correlated with reduced ECM incidence (**Extended Data Fig. 2e**). Following, we compared the durability of protection by challenging *Pb*WT and *PbP36p*^-^ RAS-immunized mice at increasing intervals post-prime (**Fig. 2c**). Both groups displayed a comparable decline in protection over time, with most mice exhibiting patent blood stage infections by 195 days (**Fig. 2d-f**), suggesting that long-term protection wanes at a similar rate regardless of parasite genotype. In contrast, *Pb*WT and *PbP36p*^-^ RAS-immunized mice that remained blood stage parasite-negative following the initial challenge 28 days after prime (**Fig. 2a,b**) maintained robust levels of sterile protection upon subsequent consecutive re-challenges at the same time points (**Extended Data Fig. 2f-i**). Taken together, these observations show that immunization with RAS exhibiting reduced infectivity but preserved CT motility can induce sterile protection with comparable durability to their wild type counterparts, reinforcing that CT is not only required (**Fig. 1**), but sufficient for effective induction of immunity by W-SPZ vaccination.

**Fig. 2.**
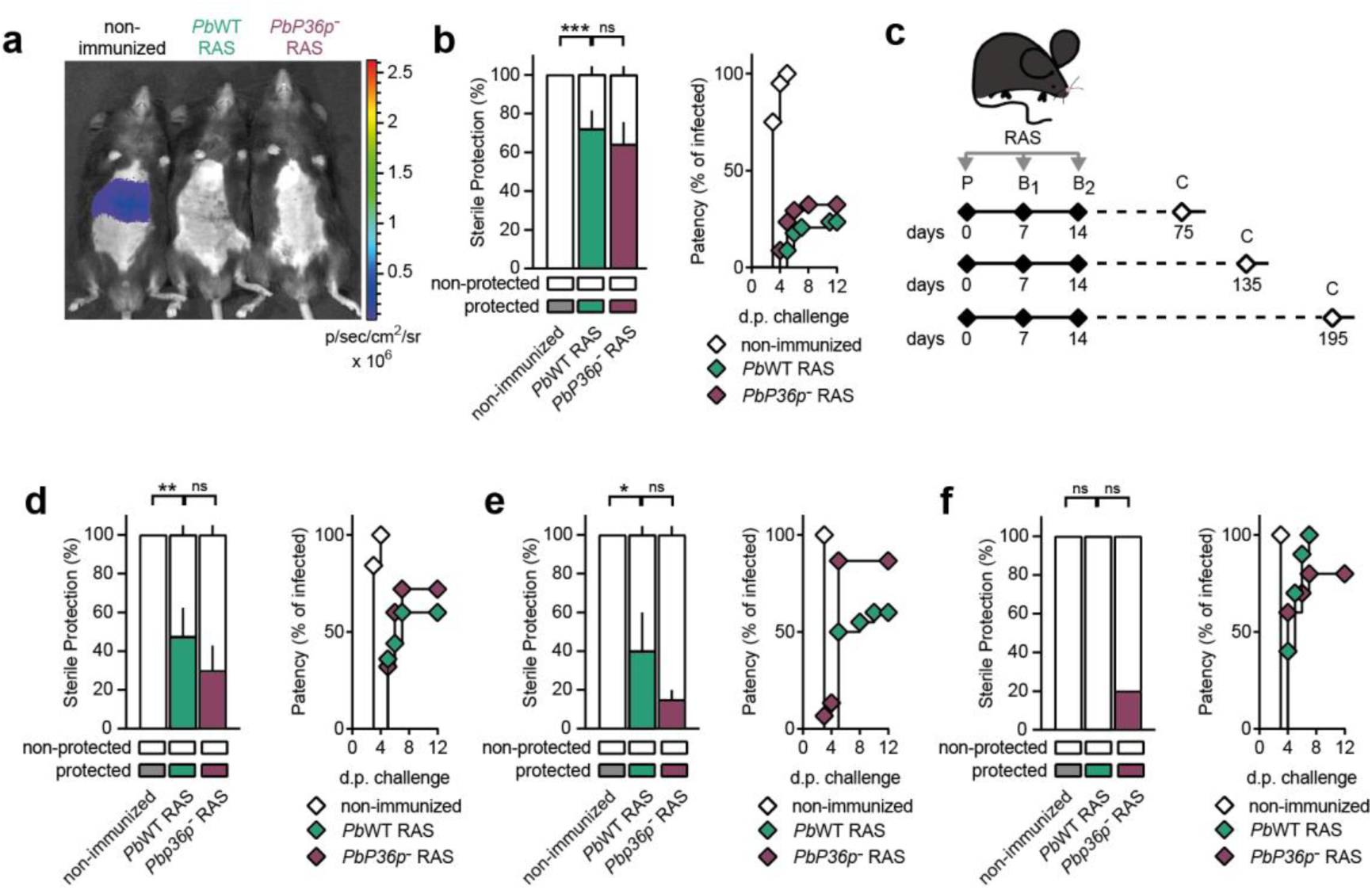
Cell traversal motility of immunizing parasites is sufficient for the establishment of sterile protection following W-SPZ RAS vaccination. **a**, Representative luciferase bioluminescence image of non-immunized, and *Pb*WT or *PbP36p^-^* RAS-immunized C57BL/6J mice, 48 hours after challenge with *Pb*WT spz. **b**, Sterile protection, represented as percentage of blood stage parasite-free mice (left), and time to patency (right) of non-immunized (n=20), and *Pb*WT (n=34) or *PbP36p^-^*(n=34) RAS-immunized C57BL/6 mice in the 12-days period following challenge with *Pb*WT spz (4-5 independent experiments). **c**, Protocol of RAS immunizations and challenge used to evaluate long-term protection. **d**-**f** Sterile protection, represented as percentage of blood stage parasite-free mice (left), and time to patency (right) of non-immunized, and *Pb*WT or *PbP36p^-^* RAS-immunized C57BL/6 mice in the 12-days period following challenge with *Pb*WT spz (**d**) 75 (n=19, n=25 and n=25, respectively) (**e**) 135 (n=12, n=20 and n=15, respectively) and (**f**) 195 (n=5, n=10 and n=10, respectively) days after prime (1-4 independent experiments). **b**, **d**, **e** and **f**; left panels, Data (Mann-Whitney test) are represented as bars ± SEM. **b**, **d**, **e** and **f**; right panels, Data (Mann-Whitney test) are represented as the mean day to patency. ns – non-significant; * - p<0.05; ** - p<0.01; *** - p<0.001. For complete statistical analysis, please refer to Source Data Fig. 2.

### Lack of cell traversal by spz during immunization hinders cellular immunity

The presence of blood stage parasites following infectious challenge of mice immunized with CT-deficient spz (**Fig. 1**) likely reflects the role of CT in promoting and/or maintaining immune mechanisms relevant for parasite clearance during hepatic infection. W-SPZ vaccination in rodents elicits both humoral and cellular responses that cooperate to eliminate spz and infected hepatocytes^26^. To investigate which of these protective immune mechanisms is hindered in the absence of CT, we assessed the dynamics of parasite elimination during challenge of *Pb*WT and *PbSpect1*^-^ RAS-immunized mice (**Fig. 3a**). At the first time point analyzed - 8 h.p.c., both RAS-immunized cohorts showed comparable reduction in parasite liver load when compared to NI controls (**Fig. 3a**), indicating early partial control of infection. Aligned with the notion that elevated anti-plasmodial antibody titers following immunization hinder parasite establishment^27,28^, both *Pb*WT and *PbSpect1*^-^ RAS-immunized mice exhibited increased, but comparable, levels of αspz IgG (**Fig. 3b**). This was also the case for *PbPlp1*^-^ and *PbP36p*^-^ RAS-immunized mice (**Extended Data Fig. 3a,b**), which is consistent with preserved antibody responses despite impaired CT or infectivity. Assessment of the temporal dynamics of infectious spz development from 16 h.p.c. onwards showed a progressive increase in liver parasite burden in *PbSpect1*^-^ RAS-immunized mice, indicative of ongoing intrahepatic development, whereas that of *Pb*WT RAS-immunized mice remained unaltered throughout infection (**Fig. 3a**). Taken together, our data show that CT is essential for the generation of protective cellular responses that eliminate infected hepatocytes, while anti-*Plasmodium* antibody production does not require either CT motility or, notably, infectivity of immunizing parasites.

**Fig. 3.**
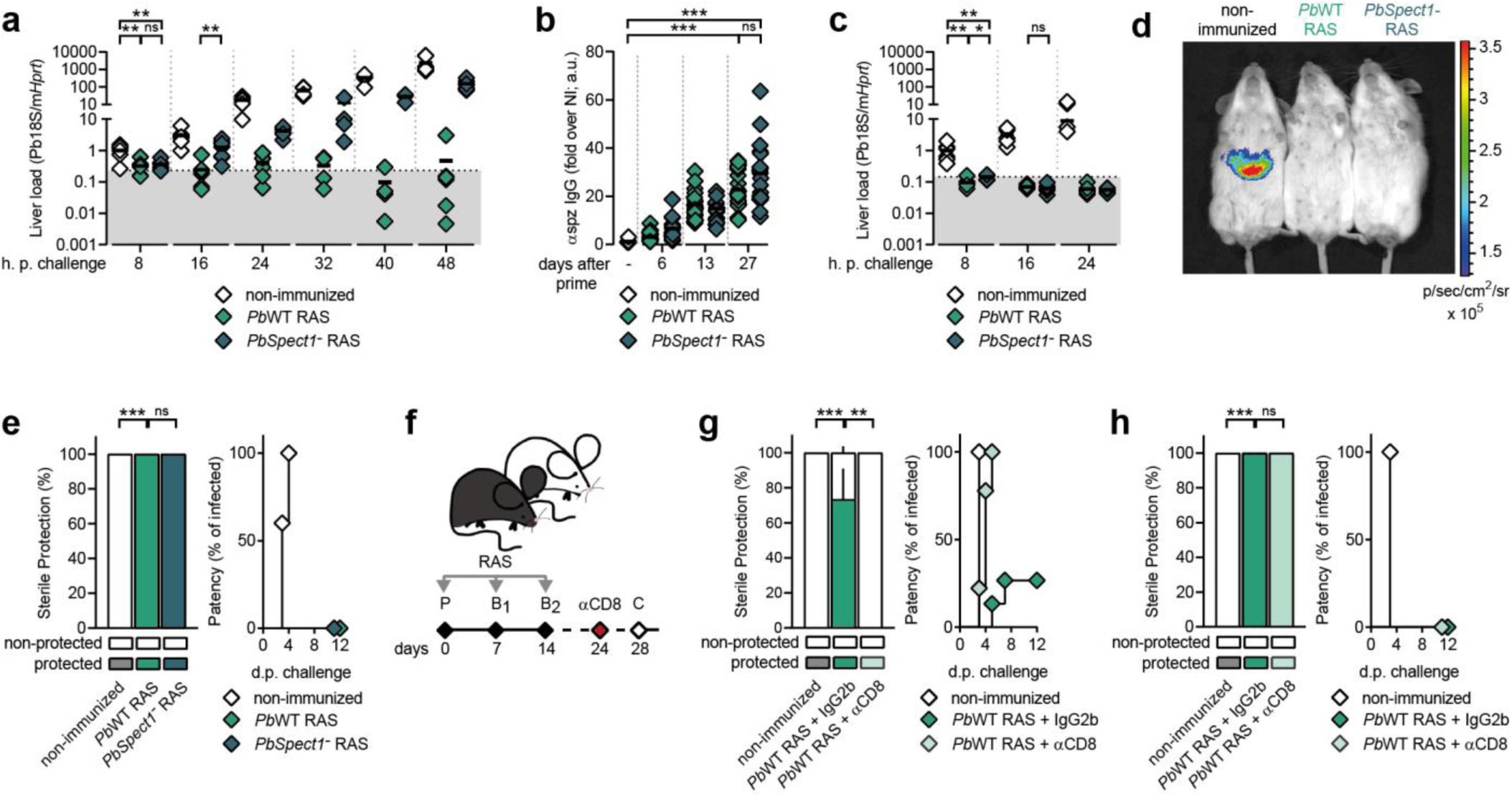
Cellular immunity, but not humoral immunity, is hindered by the lack of host cell traversal during immunization. **a**, Time-dependent quantification of *Plasmodium 18s* rRNA levels in the livers of non-immunized, and *Pb*WT or *PbSpect1^-^* RAS-immunized C57BL/6J mice at 8 (n=6), 16 (n=7), 24 (n=7), 32 (n=4), 40 (n=4), and 48 (n=8) hours after challenge with *Pb*WT spz. Gray area represents *18s* rRNA unspecific amplification signal in non-infected controls (n=6) (1-2 independent experiments). **b**, Quantification of circulating αspz IgG levels in non-immunized (n=12), and *Pb*WT (n=18) or *PbSpect1^-^* (n=18) RAS-immunized C57BL/6J mice at the indicated days after priming (2 independent experiments). **c**, Time-dependent quantification of *Plasmodium 18s* rRNA levels in the livers of non-immunized, and *Pb*WT or *PbSpect1^-^* RAS-immunized BALB/c mice at 8 (n=6), 16 (n=6) and 24 (n=6) hours after challenge with *Pb*WT spz. Gray area represents unspecific *18s* rRNA amplification signal in non-infected controls (n=7) (2 independent experiments). **d**, Representative luciferase bioluminescence image of non-immunized, and *Pb*WT or *PbSpect1^-^* RAS-immunized BALB/c mice, 48 hours after challenge with *Pb*WT spz. **e**, Sterile protection, represented as percentage of blood stage parasite-free mice (left), and time to patency (right) of non-immunized (n=10), and *Pb*WT (n=10) or *PbSpect1^-^* (n=10) RAS-immunized BALB/c mice in the 12-days period following challenge with *Pb*WT spz (2 independent experiments). **f**, Protocol of RAS immunizations and challenge following the administration of CD8 T cell-depleting antibodies. **g**, **h** Sterile protection, represented as percentage of blood stage parasite-free mice (left), and time to patency (right) of non-immunized, and *Pb*WT RAS-immunized (**g**) C57BL/6J (n=10, n=15 and n=8, respectively) or (**h**) BALB/c (n=5, n=10 and n=8, respectively) mice receiving IgG2b or αCD8 T cell-depleting antibodies followed by challenge with *Pb*WT spz (2-3 independent experiments). **a**-**c**, Data (Mann-Whitney test) are represented as dot plots, black line represents mean. **e**, **g** and **h**; left panels, Data (Mann-Whitney test) are represented as bars ± SEM. **e**, **g** and **h**; right panels, Data (Mann-Whitney test) are represented as the mean day to patency. ns – non-significant; * - p<0.05; ** - p<0.01; *** - p<0.001. For complete statistical analysis, please refer to Source Data Fig. 3.

To determine whether the requirement for CT in promoting sterile protection is specific to cellular immunity–dominated responses, we employed our RAS immunization protocol (**Fig. 1c**) in BALB/c mice, a strain known to mount strong antibody-mediated responses^29–31^. As for C57BL/6 mice (**Fig. 1b**), the 15:1 *PbSpect1*^-^ to *Pb*WT inoculum ratio yielded equivalent RAS infections at 16 h.p.i. (**Extended Data Fig. 3c**). However, analysis of the temporal dynamics of infectious parasite elimination revealed that, unlike C57BL/6 mice (**Fig. 3a**), neither *Pb*WT nor *PbSpect1*^-^RAS-immunized BALB/c mice displayed detectable liver parasite loads at any of the time points post-challenge analyzed (**Fig. 3c**). The rapid clearance of infectious spz in *Pb*WT RAS-immunized BALB/c mice correlated with the production of αspz IgG (**Extended Data Fig. 3d**), potentially leading to efficient spz neutralization prior to hepatocyte invasion^28^. Accordingly, no bioluminescence was detected in the livers of BALB/c mice immunized with either CT-competent or CT-deficient parasites (**Fig. 3d** and **Extended Data 3e**), with 100% of mice remaining blood stage parasite-free over the 12-day monitoring period (**Fig. 3e**). Together, these data suggest that CT is dispensable when sterile protection following W-SPZ immunization relies on antibody-mediated clearance of infectious parasites. To confirm that the contrasting strain-specific dependence on CT for protection (**Fig. 1d,e** and **3d,e**) reflected a dependency on cellular *versus* antibody-mediated immunity, we depleted CD8 T cells in *Pb*WT RAS-immunized C57BL/6 and BALB/c mice, 4 days prior to challenge (**Fig. 3f**). CD8 T cell depletion in C57BL/6 mice (**Extended Data Fig. 3f)** abrogated sterile protection (**Fig. 3g**), consistent with a dominant role for T cell–mediated clearance of infected hepatocytes^32,33^. In contrast, BALB/c mice remained blood stage parasite-free following CD8 T cell depletion prior to challenge (**Fig. 3h** and **Extended Data Fig. 3f**), despite the sustained reduction in CD8 T cell frequencies in the liver and the spleen throughout the 12-days monitoring period after challenge (**Extended Data Fig. 3g,h**).

### Cell traversal promotes the differentiation of CD8 effector into tissue-resident memory T cells

Having established its critical role in eliciting protective cell-mediated immunity against *Plasmodium* intrahepatic forms (**Fig. 3**), we next sought to disclose how CT shaped the hepatic immune landscape following vaccination. To this end, we employed spectral flow cytometry to profile discrete liver-infiltrating leukocyte populations in *Pb*WT and *PbSpect1^-^* RAS-immunized C57BL/6 mice, 28 days after prime. Hierarchical, unsupervised clustering of hallmark markers using FlowSOM on concatenated CD3^+^ lymphocytes and CD3^-^ cells expressing CD11b and/or NK1.1 identified 14 and 7 distinct cell clusters, respectively (**Fig. 4a** and **Extended Data Fig. 4a**). Of these, *PbSpect1*⁻ RAS-immunized mice exhibited increased frequencies of natural killer (NK) T cell clusters and of hepatic CD11b^-^ NK1.1^+^ cells, comparatively to mice immunized with *Pb*WT RAS (**Fig. 4b** and **Extended Data Fig. 4b**). NK and NK T cells have previously been associated with the elimination of *Plasmodium* pre-erythrocytic forms^34,35^, arguing against a causal link between the expansion of these cell clusters and the loss of protection observed following *PbSpect1*⁻ RAS immunizations (**Fig. 1**). In contrast, only one cluster showed a cell frequency change associated with protection. Tissue-resident memory (T_rm_) CD8 T cells were enriched in protected mice immunized with *Pb*WT RAS, but significantly under-represented in the unprotected *PbSpect1*^-^ RAS-immunized cohort (**Fig. 4b** and **Extended Data Fig. 4c**). Targeted phenotypic analysis of hepatic CD8 T cells further confirmed a significant reduction in KLRG1⁻ CD69⁺ CD8 T_rm_ cells in the livers of *PbSpect1*⁻ RAS-immunized mice (**Fig. 4c,d**). These findings suggest that CT promotes the generation of CD8 T_rm_ cells, a population required for durable and localized immunity against liver stage parasites following W-SPZ vaccination^32^. Notably, we also observed a reduction in a CD8 T cell population with an intermediate phenotype between T_eff_ and T_rm_ cells, characterized by the expression of CD69 and intermediate levels of KLRG1 (CD69^+^ KLRG1^int^), in mice immunized with *PbSpect1*⁻ RAS (**Fig. 4c,d**). It has been shown that KLRG1⁺ CD8 T_eff_ cells can downregulate KLRG1 expression and transition into long-lived memory populations - including circulating and tissue-resident subsets - while retaining high proliferative capacity and cytotoxic potential^36,37^. Given that overall CD8 T_eff_ cell numbers were comparable regardless of the genotype of immunizing spz (**Fig. 4c,d**), we hypothesized that CT promotes the conversion of CD8 T_eff_ into T_rm_ cells, with the CD69^+^ KLRG1^int^ subset representing a transitional state. To assess this, we adoptively transferred CD8 T cells isolated from *Klrg1^Cre^Rosa26^tdTomato^*reporter mice, in which KLRG1-expressing cells become irreversibly tdTomato⁺, allowing for lineage tracing even after downregulation of KLRG1^36^, into naïve C57BL/6 mice. Twenty-four hours later, recipient mice were immunized with *Pb*WT or *PbSpect1⁻* RAS, and liver and spleen leukocytes analyzed by spectral flow cytometry, 28 days post-prime (**Fig. 4e**). While no tdTomato^+^ CD8 T cells were found in the livers of non-immunized mice at the time of analysis, they constituted 58.01 ± 8.3% of liver-infiltrating *Klrg1^Cre^Rosa26^tdTomato^*CD8 T cells transferred into *Pb*WT RAS-immunized mice, with minimal presence in the spleen (**Fig. 4f**). In contrast, *PbSpect1⁻* RAS-immunized mice had reduced frequency of liver-infiltrating tdTomato⁺ cells, when compared to *Pb*WT RAS-immunized mice (**Fig. 4f**). Further phenotypic characterization of transferred CD8 T cells into *Pb*WT-immunized mice confirmed their T_rm_ cell phenotype, as 66.1 ± 8.2% of tdTomato⁺ CD8 T cells were CD69^+^, 46.08 ± 7.2% of which no longer expressed KLRG1, and exhibited cytotoxic activity following *ex vivo* stimulation (**Fig. 4g**). These findings show that CT promotes the differentiation of KLRG1⁺ CD8 T_eff_ cells into cytotoxic liver-resident memory cells.

**Fig. 4.**
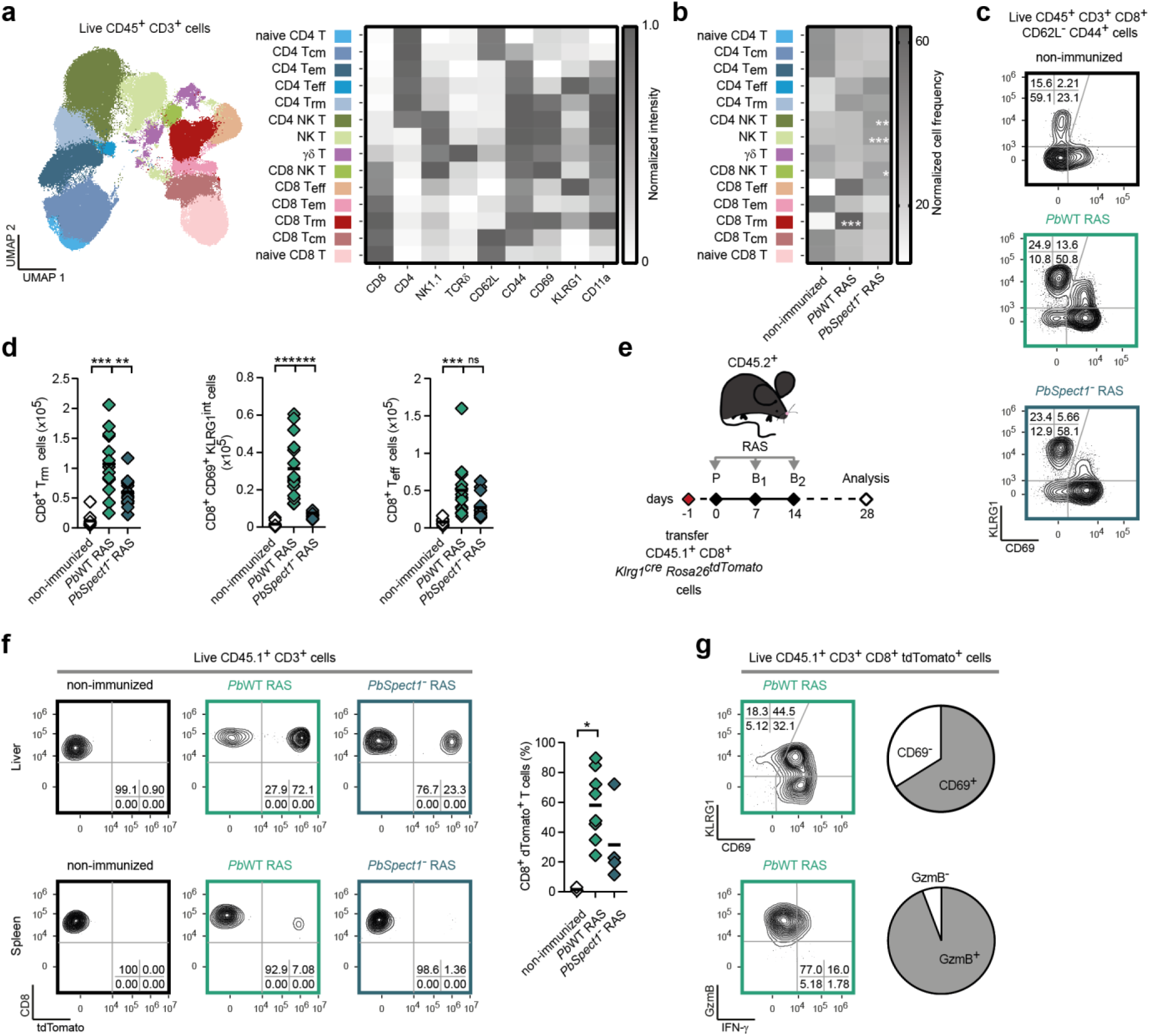
Cell traversal impacts the establishment of hepatic-resident memory CD8 T cells following W-SPZ vaccination. **a**, Uniform manifold approximation and projection (UMAP) representation (left) and mean fluorescence intensity (MFI) of each hallmark marker (right) in immune cell clusters identified by FlowSOM in concatenated CD3^+^ immune cells isolated from the livers of non-infected (n=6), and *Pb*WT (n=10) or *PbSpect1^-^* (n=10) RAS-immunized C57BL/6J mice, 28 days after prime (2 independent experiments). T_cm_ – central memory T cell; T_em_ – effector memory T cell; T_eff_ – effector T cell; T_rm_ – tissue-resident memory T cell; NK – natural killer. Cell gating strategy shown in **Extended Data Fig. 5a. b**, Heatmap of normalized cell frequencies within each identified cluster of CD3^+^ lymphocytes in the livers of non-infected, and *Pb*WT or *PbSpect1^-^* RAS-immunized C57BL/6J mice, 28 days after prime. **c, d** Representative flow cytometry plots (**c**) and quantification (**d**) of tissue-resident memory (T_rm_) (left), CD69^+^ KLRG^int^ (middle) and effector (T_eff_) (right) CD8 T cells in the livers of non-immunized (n=9), and *Pb*WT (n=15) or *PbSpect1^-^* (n=15) RAS-immunized C57BL/6J mice, 28 days after prime (3 independent experiments). Cell gating strategy shown in **Extended Data Fig. 5b. e**, Protocol of adoptive transfer of KLRG1 reporter CD8 T cells and RAS immunizations. **f**, Representative flow cytometry plots (left) and frequency (right) of tdTomato-positive, transferred KLRG1 reporter CD8 T cells in the livers and spleens of non-immunized (n=3), and *Pb*WT (n=8) or *PbSpect1^-^* (n=4) RAS-immunized C57BL/6J mice 28 days after prime (1 to 2 independent experiments). Cell gating strategy shown in **Extended Data Fig. 5b. g**, Representative flow cytometry plots and frequency of CD69 (left) and granzyme B (right) expression in tdTomato-positive, transferred KLRG1 reporter CD8 T cells in the livers of *Pb*WT RAS-immunized C57BL/6J mice (n=8), 28 days after prime (2 independent experiments). Cell gating strategy shown in **Extended Data Fig. 5b. b**, Data (Mann-Whitney test) are represented as heatmap. **d**, **f** Data (Mann-Whitney test) are represented as dot plots, black line represents mean. **g**, Data are represented as pie charts. ns – non-significant; * - p<0.05; ** - p<0.01; *** - p<0.001. For complete statistical analysis, please refer to Source Data Fig. 4.

## DISCUSSION

Despite major advances in our understanding of host immune responses during *Plasmodium* liver infection, parasite-intrinsic determinants that shape these responses remain poorly defined.

Efforts in vaccine development have mostly focused on identifying and targeting specific antigens or epitopes capable of eliciting protective immune responses. Our findings challenge this antigen-centric view by demonstrating that parasite behavior traits are a critical contributor to cellular immunity elicited during W-SPZ immunization.

In this study, we showed that CT by immunizing parasites, which occurs before hepatocyte invasion^4^, was both required and sufficient for inducing sterilizing immunity following W-SPZ vaccination. This finding contrasts with the prevailing view that vaccine-induced protection is primarily driven by productive hepatocyte infection and by the extent of intrahepatic parasite development^38^. Using multiple genetic parasite models, we showed that even when CT-deficient parasites were allowed to fully replicate in hepatocytes, they failed to confer protection. Conversely, CT-competent parasites that are replication-impaired still elicited robust protection, equivalent to wild-type parasites. These data establish CT as a key immunogenic event and uncouples sterilizing protection following W-SPZ vaccination from productive *Plasmodium* infection.

RAS-based immunizations elicit both humoral and cellular responses, each contributing to preclude re-infection of immunized hosts^32,39^. The observation that sterile protection was abrogated in C57BL/6 mice immunized with CT-deficient parasites, despite high αspz antibody levels, underscores that CT contributes to cell-mediated, but not antibody-mediated, immunity. Consistent with this and aligning with observations by others^40^, CD8 T cell depletion in CT-competent RAS-immunized C57BL/6 mice abrogated protection. In contrast, in BALB/c mice, which favor antibody-mediated immunity^29–31^, both CT-competent and -deficient immunizing parasites elicited comparable levels of protection, independent of CD8 T cells. These results highlight how host genetics dictates immune correlates of protection, and underscore CT as a non-redundant component for inducing CD8 T cell–mediated immunity in hosts where this response dominates.

Cellular responses, particularly those dependent on CD8 T cells, are essential for the clearance of infected hepatocytes,^40^ with prolonged protection following malaria vaccination being associated with the presence of liver-resident memory CD8 T (T_rm_) cells^32^. Using unsupervised clustering of liver leukocytes, we found that CD8 T_rm_ cells expanded significantly following immunization with CT-competent *versus* CT-deficient RAS. Within these, a subset of CD8 T cells phenotypically characterized by the intermediate expression of the activation-associated molecule KLRG1 (CD69⁺ KLRG1^int^) was highly reduced in CT-deficient immunized mice that failed to develop protection. This specific T_rm_ subset has been previously shown to expand following RAS vaccination^41^, while its reduced frequency associates with low vaccine efficacy^12^. Now, we show that CT by immunizing parasites drives the conversion of CD8 T_eff_ cells into T_rm_ cells, linking a parasite behavioral trait to the establishment of a protective memory T cell subset. The mechanism by which CT promotes T_rm_ differentiation remains to be defined. However, environmental factors, including inflammatory cues, promote T_rm_ cell formation^33,42,43^. CT involves the breaching of host cell membranes, which can result in the release of intracellular molecules that act as danger-associated molecular patterns^44^ or trigger transcriptional stress responses by wounding cells^45^, both of which may contribute to a pro-inflammatory hepatic microenvironment conducive to T_rm_ cell differentiation. While the precise contribution of these cells to the elimination of infected hepatocytes has yet to be elucidated, their presence in mice immunized with CT-competent parasites, which exhibit robust protection, contrasts with their absence in non-protected *PbSpect1⁻* RAS-immunized hosts. Such association supports the notion that enrichment of hepatic T_rm_ cell populations via conversion of CD8 T_eff_ cells in a CT-dependent manner contributes to sterilizing immunity against *Plasmodium* liver infection.

These findings carry important implications for the development of malaria vaccines for human use. Traditionally, vaccine design has prioritized the search for optimal antigens, epitopes, and immune signatures. While this approach remains essential, our findings linking CT and protective immunity should broaden the focus from solely considering the antigenic content of immunizing strategies to include the dynamic interactions between the parasite and host tissues. This conceptual shift opens new avenues for malaria vaccine development, where parasite behavior can be harnessed to shape effective and durable immunity. Responses to W-SPZ vaccines in humans are known to be heterogeneous, with a fraction of vaccinees failing to develop sterile protection^10,11^. The reasons behind such variability are not fully understood but may include differences in vaccine delivery, host genetics, and immune history. Our data raise the possibility that parasite behavior during vaccination, such as its capacity to traverse host cells, could be a previously unrecognized contributor to such heterogeneity. Next-generation W-SPZ vaccines may benefit from engineering or selecting immunizing parasites with optimized behavioral properties, particularly enhanced CT motility, to promote effective and long-lasting CD8 T_rm_ cell responses. Moreover, the currently licensed malaria vaccines RTS,S and R21, consisting of subunit formulations targeting the circumsporozoite protein of *P. falciparum*, rely heavily on humoral immunity^46^ and consistently fail to prevent infection, albeit reducing clinical disease burden^6–9^. The identification of molecular and inflammatory cues induced by CT may enable the design of adjuvants that replicate its immunostimulatory effect in cellular immunity, extending the applicability of this principle beyond W-SPZ vaccines.

## Acknowledgments

We are thankful to: Takaharu Okada (RIKEN Center for Integrative Medical Sciences, Japan) for providing the C57BL/6 *Klgr1^Cre^Rosa26^tdTomato^* reporter mouse strain; Ana Parreira (GIMM, Portugal) for mosquito rearing and infection with *Plasmodium* parasites; Inês Bento and Karine Serre for critically reviewing this manuscript, and the Flow Cytometry and Rodent Facilities teams (GIMM, Portugal) for assistance. This work was supported by the European Commission (101097801-PASSAGE ERC-2022-ADG) and Fundação para a Ciência e Tecnologia, Portugal (PTDC/MED-IMU/28664/2017), attributed to M.M.M. and Â.F.C., respectively. A.R. was supported by the European Commission (101097801-PASSAGE ERC-2022-ADG). H.N.C. was supported by the “la Caixa” Banking Foundation (HR21-00848). N.V.L., C.F. and Â.F.C. were supported by Fundação para a Ciência e Tecnologia, Portugal (2022.04143.PTDC, 2021.01136.CEECIND and 2023.06409.CEECIND, respectively).

## Author contributions

Conceptualization, M.M.M. and Â.F.C.; methodology, A.R. and Â.F.C.; investigation, A.R., A.M.M., H.N.C., S.M., N.V.L., C.F. and Â.F.C.; formal analysis, A.R. and R.G.; resources, M.V. and M.P.; writing– original draft, A.R., M.M.M. and Â.F.C.; writing– review & editing, M.V. and M.P.; visualization, A.R. and Â.F.C.; funding acquisition, M.M.M. and Â.F.C.

## Competing interests

The authors declare no competing interests.

## Materials availability

This study generated a new *Plasmodium berghei* line genetically deficient for the *perforin-like protein 1* gene that is available upon request.

## Data availability

All data are available in the main text and in the source data, and can be assessed at

---

**Extended Data Fig. 1.**
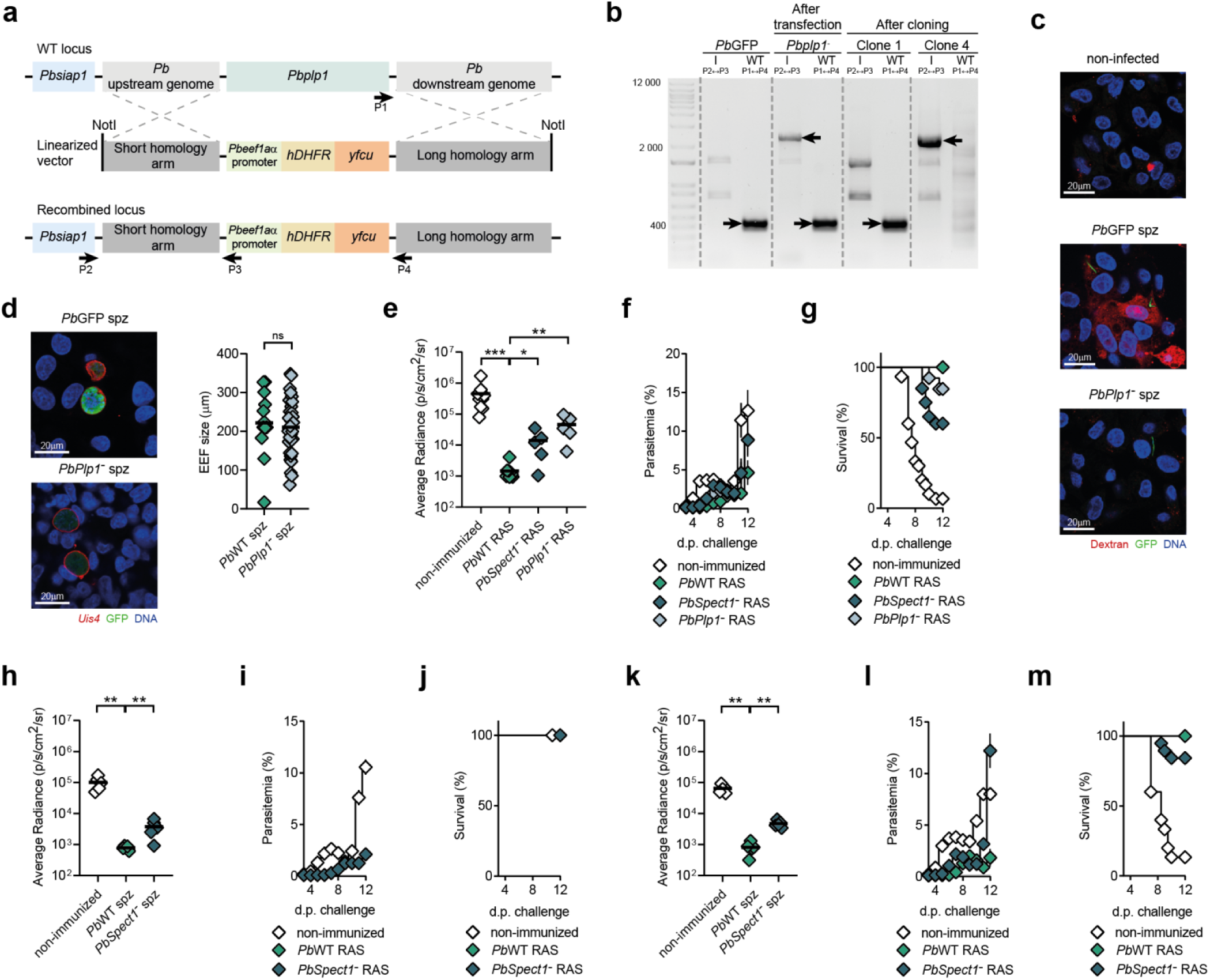
Generation and characterization of the *PbPlp1*^-^ parasite line and clinical outcome following W-SPZ vaccination and challenge. **a**, Schematic of the strategy followed to generate the *PbPlp1*^-^ parasite line, representing the PBANKA_100630 WT locus (top), the PbGEM 293440 linearized vector (middle), and the recombined locus (bottom). Dashed lines represent homologous recombination sites for the integration of the *Pb*GEM-293340 vector into the *Pb*WT genome. The recombined locus contains a *Pb* elongation factor 1-alpha (*PbEef1aα*) promoter, followed by a resistance cassette composed of the human dihydrofolate reductase (*hDHFR*) gene and the bifunctional yeast enzyme cytosine deaminase and uridyl phosphoribosyl transferase (*yfcu*). Arrows represent the annealing site of primers used for parasite genotyping. **b**, Electrophoretic analysis of *PbPlp1*^-^ parasites after cloning. Bands corresponding to the presence of the *Plp1* wild-type locus are labeled with black arrows pointing right, and bands corresponding to the correct integration of the resistance cassette are labeled with black arrows pointing left. **c**, Representative fluorescence confocal microscopy images of HepG2 cells either non-infected (left) or infected, 2 hours earlier with *Pb*GFP (middle) or *PbPlp1*^-^ (right) spz in the presence of rhodamine dextran. GFP labels sporozoites; rhodamin dextran indicates traversed cells, and Hoescht 33342 stains DNA (2 independent experiments). Images were acquired with 63x magnification and scale bar represents 20 µm. **d**, Representative fluorescence confocal microscopy images (left) and quantification of EEF size (right) of *Pb*GFP or *PbPlp1*^-^ parasites, 48 hours post infection of HepG2 cells. UIS4 labels the parasitophorous vacuole membrane; GFP labels parasite cytoplasm, and Hoescht 33342 stains DNA (3 independent experiments). Images were acquired with 63x magnification and scale bar represents 20 µm. **e**, Quantification of functional luciferase activity in non-immunized (n=5), and *Pb*WT (n=5), *PbSpect1^-^* (n=5) or *PbPlp1^-^* (n=5) RAS-immunized C57BL/6J mice, 48 hours after challenge with *Pb*WT spz. **f, g** Parasitemia (**f**) and survival (**g**) in non-immunized (n=35), and patent *Pb*WT (n=40), *PbSpect1^-^* (n=20) or *PbPlp1^-^* (n=15) RAS-immunized C57BL/6J mice in the 12-days period following challenge with *Pb*WT spz (3-8 independent experiments). Parasitemia gating strategy shown in **Extended Data** Fig. 5c. **h**, Quantification of functional luciferase activity in non-immunized (n=5), and *Pb*WT (n=5) or *PbSpect1^-^* (n=5) CPS-AZ-immunized C57BL/6 mice, 48 hours after challenge with *Pb*WT spz. **i, j** Parasitemia (**i**) and survival (**j**) in non-immunized (n=5), and patent *Pb*WT (n=15) or *PbSpect1^-^* (n=10) CPS-AZ-immunized C57BL/6 mice in the 12-days period following challenge with *Pb*WT spz (1-3 independent experiments). Parasitemia gating strategy shown in **Extended Data** Fig. 5c. **k**, Quantification of functional luciferase activity in non-immunized (n=5), and *Pb*WT (n=5) or *PbSpect1^-^* (n=5) CPS-CQ-immunized C57BL/6 mice, 48 hours after challenge with *Pb*WT spz. **l**, **m** Parasitemia (**l**) and survival (**m**) in non-immunized (n=15), and patent *Pb*WT (n=25) or *PbSpect1^-^* (n=19) CPS-CQ-immunized C57BL/6 mice in the 12-days period following challenge with *Pb*WT spz (3-4 independent experiments). Parasitemia gating strategy shown in **Extended Data** Fig. 5c. **d**, **e**, **h** and **k** Data (Mann-Whitney test) are represented as dot plots, black line represents mean. **f**, **i** and **l** Data (Mann-Whitney test) are represented as mean percentage of infected erythrocytes ± SEM. **g**, **j** and **m** Data (Log rank (Mantel-Cox) test) are represented as mean survival rate. ns – non-significant; * - p<0.05; ** - p<0.01; *** - p<0.001. For complete statistical analysis, please refer to Source Data Extended Data Fig.1.

**Extended Data Fig. 2.**
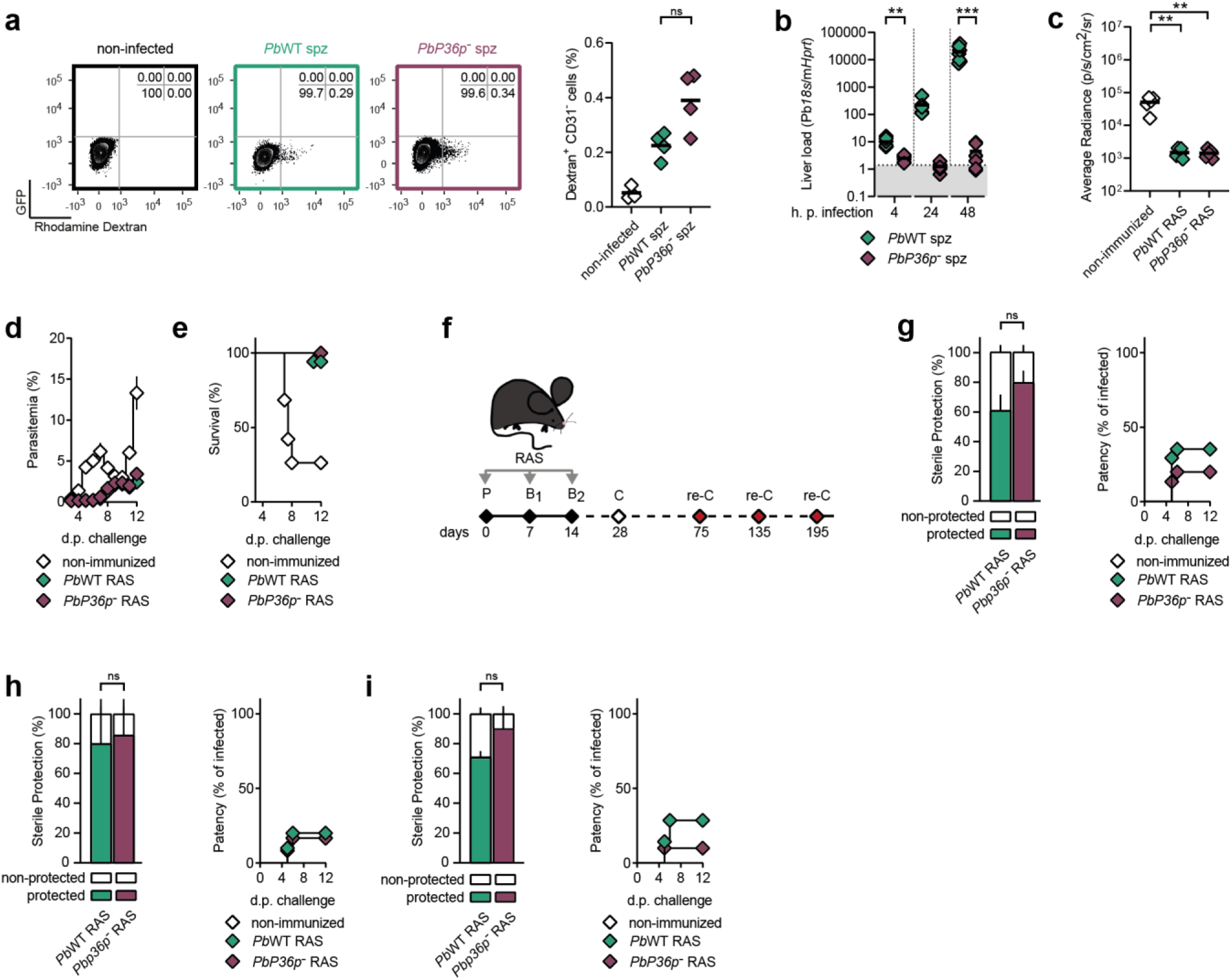
In vivo characterization of *PbP36p^-^* parasite line and sequential challenge protocol. **a**, Representative flow cytometry plots and frequency of traversed, rhodamine dextran-positive liver parenchymal cells in the livers of C57BL/6 mice, either non-infected (n=3), or 2 hours after receiving 5x10^5^ *Pb*WT (n=4) or 5x10^5^ *PbP36p^-^*(n=4) spz (3 independent experiments). Cell gating strategy shown in **Extended Data Fig. 5d. b**, Time-dependent quantification of *Plasmodium 18s* rRNA levels in the livers of C57BL/6 mice receiving 10^4^ *Pb*WT or *PbP36p^-^* spz at 4 (n=6 per group), 24 (n=6 and n=8, respectively) and 48 (n=6 and n=8, respectively) hours after infection (2 independent experiments). Gray area represents unspecific *18s* rRNA amplification signal in non-infected controls (n=5). **c**, Quantification of functional luciferase activity in non-immunized, and *Pb*WT or *PbP36p^-^* RAS-immunized C57BL/6 mice, 48 hours after challenge with *Pb*WT spz. **d**, **e** Parasitemia (**d**) and survival (**e**) in non-immunized (n=20), and patent *Pb*WT (n=34) or *PbP36p^-^* (n=34) RAS-immunized C57BL/6 mice in the 12-days period following challenge with *Pb*WT spz (4-5 independent experiments). Parasitemia gating strategy shown in **Extended Data Fig. 5c. f**, Protocol used for RAS immunizations to evaluate long-term protection following successive re-challenges. **g**-**i** Sterile protection, represented as percentage of blood stage parasite-free mice (left), and time to patency (right) of *Pb*WT or *PbP36p^-^*RAS-immunized C57BL/6 mice following re-challenge with *Pb*WT spz (**g**) 75 (n=15 and n=15, respectively) (**h**) 135 (n=9 and n=12, respectively) and (**i**) 195 (n=7 and n=10, respectively) days after prime (2 independent experiments). **a**, **b** and **c** Data (Mann-Whitney test) are represented as dot plots, black line represents mean. **d**, Data (Mann-Whitney test) are represented as mean percentage of infected erythrocytes ± SEM. **e**, Data (Log rank (Mantel-Cox) test) are represented as mean survival rate. **g**-**i**; left panels, Data (Mann-Whitney test) are represented as bars ± SEM. **g**-**i**; right panels, Data (Mann-Whitney test) are represented as the mean day to patency. ns – non-significant; * - p<0.05; ** - p<0.01; *** - p<0.001. For complete statistical analysis, please refer to Source Data Extended Data Fig.2.

**Extended Data Fig. 3.**
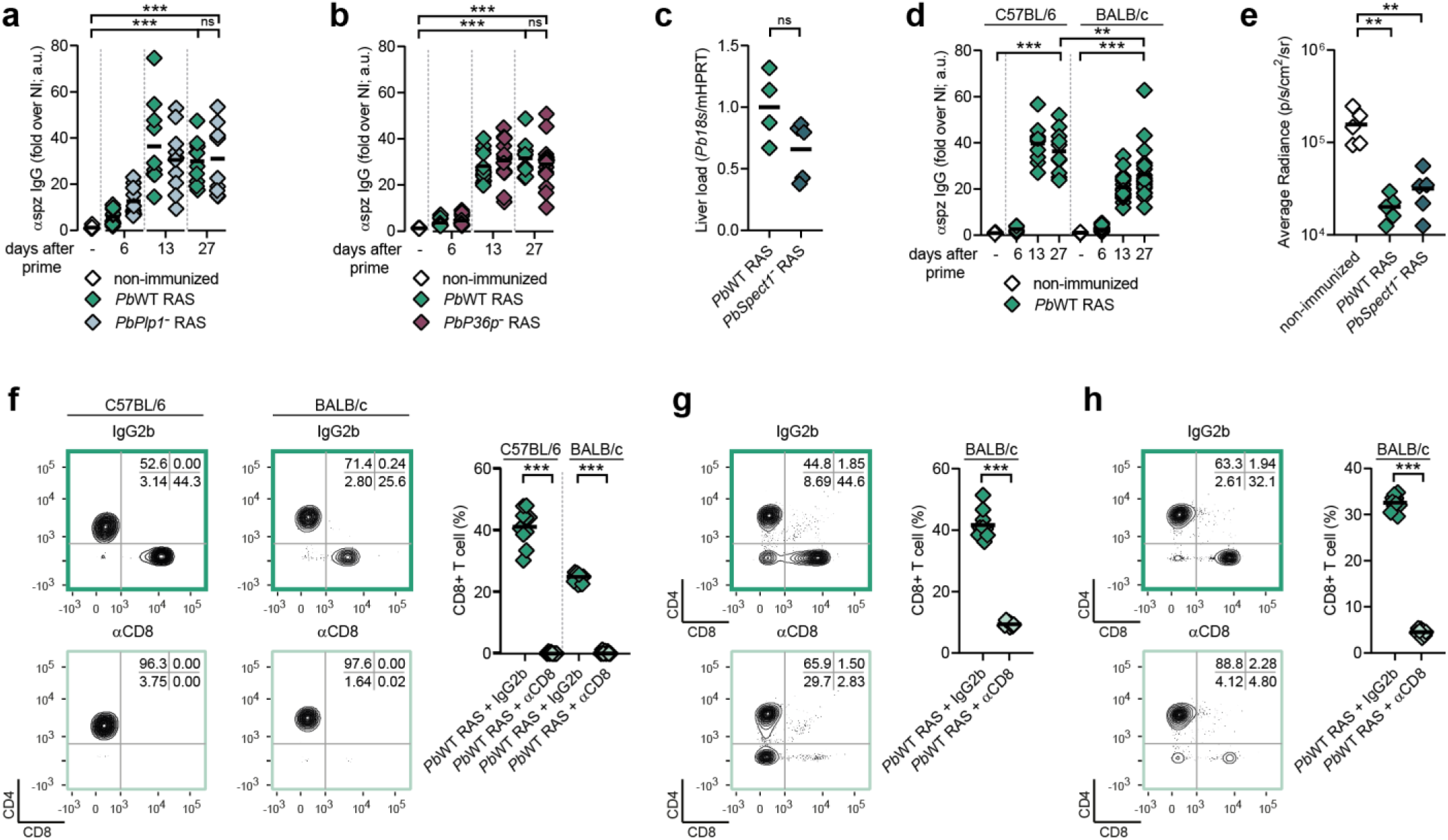
BALB/c humoral responses following W-SPZ vaccination and assessment of CD8 T cell depletion. **a**, Quantification of circulating αspz IgG levels in non-immunized (n=10), and *Pb*WT (n=10) or *PbPlp1^-^* (n=10) RAS-immunized C57BL/6 mice at the indicated days after priming (2 independent experiments). **b**, Quantification of circulating αspz IgG levels in non-immunized (n=5), and *Pb*WT (n=10-15 per time point) or *PbP36p^-^* (n=15) RAS-immunized C57BL/6J mice at the indicated days after prime (1-2 independent experiments). **c**, Quantification of *Plasmodium 18s* rRNA levels in the livers of BALB/c mice, 16 hours after inoculation of 10^4^ *Pb*WT (n=4) or 15x10^4^ *PbSpect1^-^* (n=5) RAS (1 experiment). **d**, Quantification of circulating αspz IgG levels in C57BL/6 mice, either non-immunized (n=5) or immunized with *Pb*WT RAS ( n=10 per time point), and BALB/c mice, either non-immunized (n=10) or immunized with *Pb*WT RAS (n=15-20 per time point) at the indicated days after priming (1-2 independent experiments). **e**, Quantification of functional luciferase activity in non-immunized, and *Pb*WT or *PbSpect1p^-^* RAS-immunized BALB/c mice, 48 hours after challenge with *Pb*WT spz. **f**, Representative flow cytometry plots (right) and frequency (left) of CD4 and CD8 T cells in the peripheral blood of *Pb*WT RAS-immunized C57BL/6 or BALB/c at the time of challenge after receiving, 4 days earlier, IgG2b (n=10 per group) or αCD8 T cell-depleting (n=8 per group) antibodies (2 independent experiments). Cell gating strategy shown in **Extended Data Fig. 5e. g**, **h** Representative flow cytometry plots and frequency of CD4 and CD8 T cells in (**g**) the livers and (**h**) the spleens of protected *Pb*WT RAS-immunized BALB/c mice receiving IgG2b (n=10) or αCD8 T cell-depleting (n=8) antibodies 4 days prior to challenge, analyzed at the end of the 12-day period after challenge (2 independent experiments). Cell gating strategy shown in **Extended Data Fig. 5e. a**-**h** Data (Mann-Whitney test) are represented as dot plots, black line represents mean. ns – non-significant; * - p<0.05; ** - p<0.01; *** - p<0.001. For complete statistical analysis, please refer to Source Data Extended Data Fig. 3.

**Extended Data Fig. 4.**
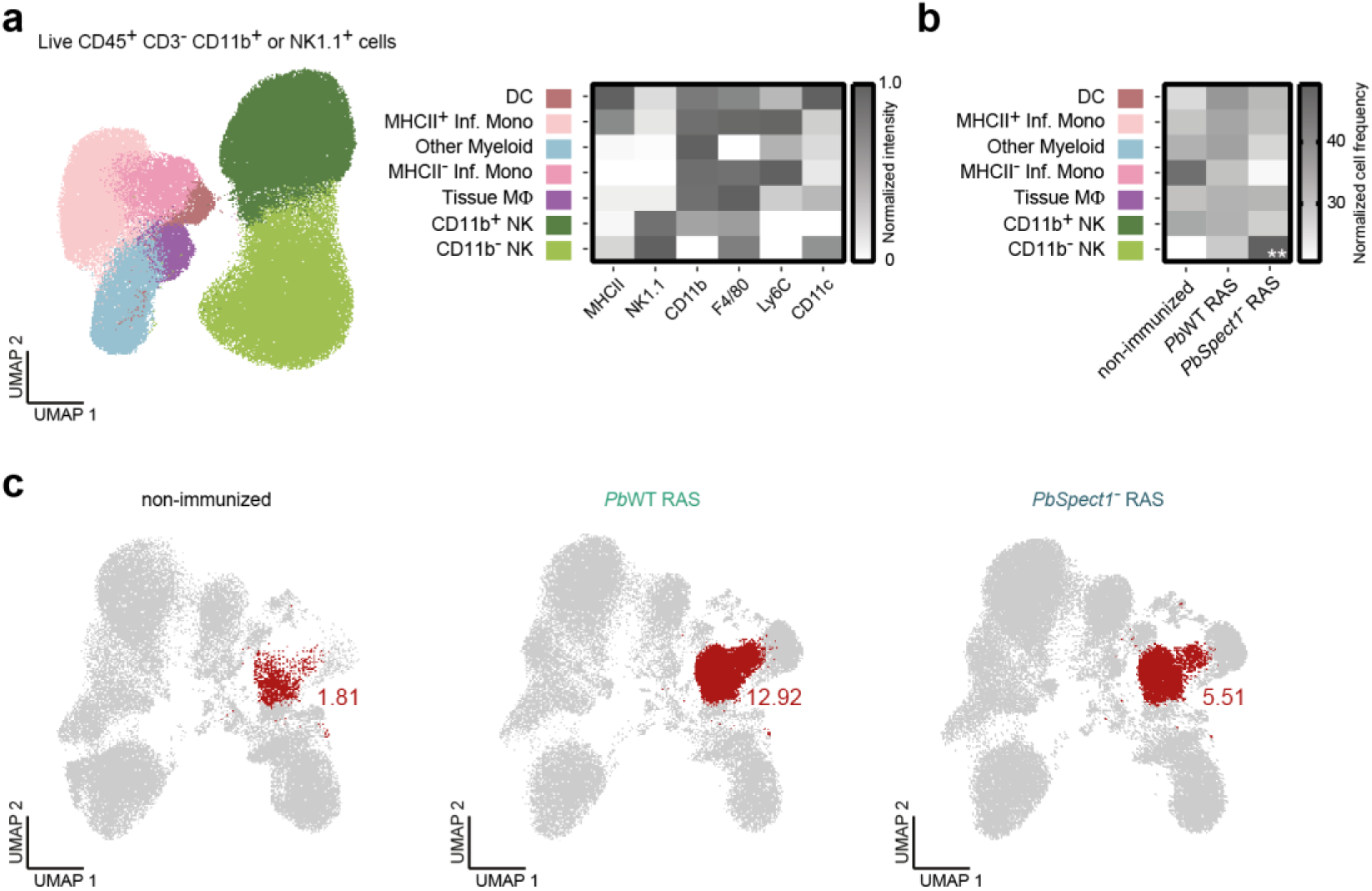
Innate and CD8 T_rm_ immune responses following W-SPZ vaccination. **a**, Uniform manifold approximation and projection (UMAP) representation (left) and mean fluorescence intensity (MFI) of each hallmark marker (right) in immune cell clusters identified by FlowSOM in concatenated CD3^-^ immune cells expressing CD11b and/or NK1.1 isolated from the livers of non-infected (n=5), and *Pb*WT (n=9) or *PbSpect1^-^* (n=9) RAS-immunized C57BL/6J mice, 28 days after prime (2 independent experiments). DC – dendritic cell; Inf. Mono – inflammatory monocyte; MΦ – macrophage; NK – natural killer. Cell gating strategy shown in **Extended Data Fig. 5a. b**, Heatmap of normalized cell frequencies within each identified cluster of CD3⁻ immune cells expressing CD11b and/or NK1.1 in the livers of non-infected, and *Pb*WT or *PbSpect1^-^* RAS-immunized C57BL/6J mice, 28 days after prime. **c**, Uniform manifold approximation and projection (UMAP) representation of concatenated CD3^+^ immune cells isolated from the livers of non-infected (n=6) (left), and PbWT (n=10) (middle) or PbSpect1- (n=10) (right) RAS-immunized C57BL/6J mice, 28 days after prime (2 independent experiments). Frequencies of the highlighted CD8 T_rm_ cell cluster (red) are shown for each condition. **b**, Data (Mann-Whitney test) are represented as a heatmap. ns – non-significant; * - p<0.05; ** - p<0.01; *** - p<0.001. For complete statistical analysis, please refer to Source Data Extended Data Fig. 4.

**Extended Data Fig. 5.**
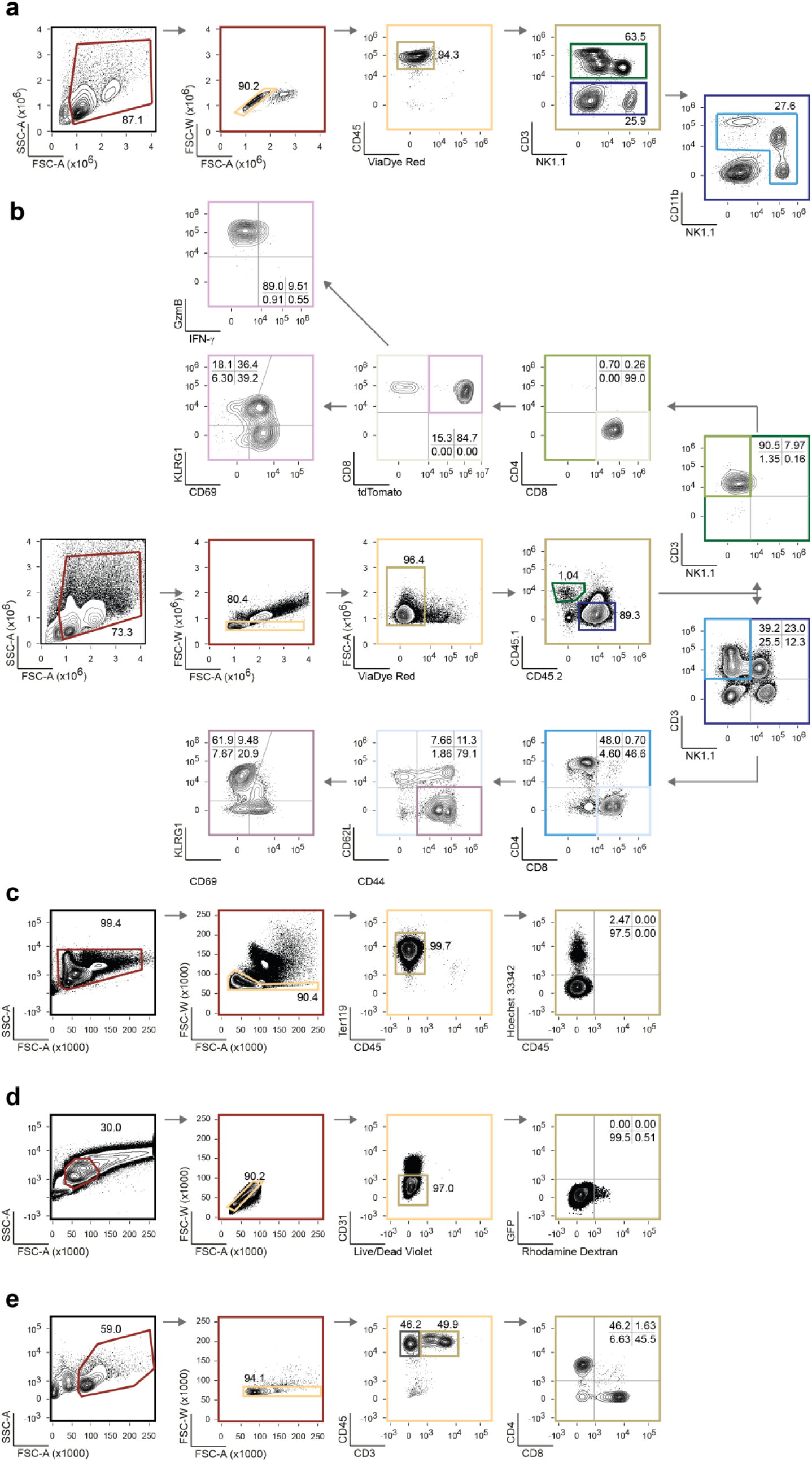
Gating strategies. **a**, Gating strategy used for downstream cell clustering of liver-isolated leukocytes. Leukocytes were gated using the forward scatter (FSC-A) and side scatter (SSC-A) parameters, cell aggregates were excluded using FSC-A *vs*. FSC-W, and live leukocytes were selected based on live/dead discrimination in a CD45 *vs*. ViaDye Red plot. Lymphocytes and natural killer T cells were selected for concatenation based on the expression of CD3 and NK1.1 in a CD3 *vs*. NK1.1 plot. CD3^-^ leukocytes were further gated based on the expression of CD11b and/or NK1.1 for concatenation and analysis of myeloid and natural killer subsets. **b**, Gating strategy used to evaluate the frequency of endogenous and transferred KLRG1 reporter effector and memory CD8 T cell subsets. Leukocytes isolated from livers or spleens were gated using the forward scatter (FSC-A) and side scatter (SSC-A) parameters, cell aggregates were excluded using FSC-A *vs*. FSC-W, and live leukocytes were selected based on live/dead discrimination in a FSC-A *vs*. ViaDye Red plot. CD45 isoforms were used to discriminate between endogenous (CD45.2) leukocytes and transferred KLRG1 reporter (CD45.1) lymphocytes. Endogenous CD8 lymphocytes were selected based on the expression of CD3 in a CD3 *vs*. NK1.1 plot and further gated based on the expression of CD8. Activated CD8 lymphocytes were discriminated in a CD44 *vs*. CD62L plot, and analyzed for the expression of KLRG1 and CD69. Transferred KLRG1 reporter lymphocytes were selected based on the expression of CD3 and CD8, and tdTomato conversion evaluated in a CD8 *vs*. tdTomato plot. tdTomato-positive CD8 lymphocytes were analyzed for the expression of KLRG1 and CD69 and for the production of granzyme B *vs*. the production of IFN-γ. **c**, Gating strategy used to analyze the frequency of infected erythrocytes (parasitemia). Erythrocytes were gated using the forward scatter (FSC-A) and side scatter (SSC-A) parameters, cell aggregates were excluded using FSC-A vs. FSC-W, and erythrocytes were selected based on Ter119 expression in a CD45 *vs.* TER119 plot. DNA content using Hoechst 33342 discriminated between infected and non-infected erythrocytes in a Hoechst 33342 *vs*. CD45 plot. **d**, Gating strategy used to analyze the frequency of traversed liver parenchyma cells. Liver parenchyma cells were gated using the forward scatter (FSC-A) and side scatter (SSC-A) parameters, cell aggregates were excluded using FSC-A *vs*. FSC-W, endothelial cells were excluded based on CD31 expression, and live parenchyma cells were selected in a CD31 *vs.* live/dead dye plot. Traversed cells were identified based on their content of rhodamine dextran in a GFP *vs*. rhodamine dextran plot. **e**, Gating strategy used to evaluate the depletion of CD8 T cells. Circulating leukocytes, or isolated from livers or spleens, were gated using the forward scatter (FSC-A) and side scatter (SSC-A) parameters, cell aggregates were excluded using FSC-A *vs*. FSC-W, and lymphocytes were selected based on CD45 and CD3 expression. Lymphocytes were further gated according to the expression of CD8 and CD4.

## METHODS

### Mice

C57BL/6 and BALB/c wild-type mice were obtained from Charles River Laboratories International Inc. *Klgr1^Cre^Rosa26^tdTomato^*mice (originally obtained from Takaharu Okada, RIKEN Center for Integrative Medical Sciences, Japan^36^), were bred at the Gulbenkian Institute for Molecular Medicine (GIMM), Lisbon, Portugal. All mice in this study were housed in specific pathogen-free conditions. For immunizations and infections, both male and female mice (5-6 weeks old) were used. For *Klgr1^Cre^Rosa26^tdTomato^* CD8 T cell transfers, male donors (14-27 weeks old) were used. Animal experimentation protocols were approved by the institutional animal welfare body – ORBEA-GIMM – and by the Direcção-Geral de Alimentação e Veterinária. All procedures were performed in strict accordance with the Directive 2010/63/EU for the care and use of animals for scientific purposes.

### Parasites

For this study, *Anopheles stephensi* female mosquitoes, bred and housed at GIMM Insectary, Lisbon, Portugal, were infected with the following *Plasmodium berghei* ANKA (*Pb*ANKA) parasite lines: (i) *Pb*ANKA expressing a GFP-Luciferase fusion protein (676m1cl1; RMgm-29)^47^, herein referred to as *Pb*WT; (ii) marker-free *Pb*ANKA expressing GFP (2.34)^48^, herein referred to as *Pb*GFP; (iii) *Pb*ANKA genetically deleted for the *Sporozoite microneme protein essential for cell traversal 1* (*Spect1*) gene (RMgm-138)^18^, herein referred to as *PbSpect1*^-^; (iv) *Pb*ANKA genetically deleted for the *P36p* gene (417m0cl2; RMgm-41)^25^, herein referred to as *PbP36p*^-^; and (v) *Pb*ANKA genetically deleted for the *perforin-like protein 1* (*Plp1*^-^) gene, generated in this study and herein referred to as *PbPlp1*^-^. The generation of *PbPlp1*^-^ was carried out using the *Pb*GFP parasite line and the *Pb*GEM-293340 (PlasmoDB) vector. *Pb*GEM-293440 contains a modified locus of the PBANKA_100630 gene (*Plp1*; PlasmoDB), consisting of the *Pbeef1aa* promoter, followed by the human dihydrofolate reductase (*hDHFR*) gene, and the bifunctional yeast enzyme cytosine deaminase and uridyl phosphoribosyl transferase (*yFCU*). The modified target locus was integrated in the *Pb*GFP parasite line by double crossover homologous recombination, recombinant parasites were selected with pyrimethamine and dilution cloning were carried out as previously described^47^. The genomic DNA (gDNA) of parasite clones was isolated from parasitized blood using the NZY Blood gDNA Isolation kit (NZYtech), and genotyped by polymerase chain reaction (PCR) using NZYTaq II 2x Colourless Master Mix (0.1U/µL; NZYtech,) and primers specific for the *Plp1* wild-type locus (5′ primer P2 (ACCGTTAGATCTGATTGCGGCCA) and 3′ primer P4 (ACACATAGCGAAACCATGTTGTCT)) and for the correct integration of the modified target locus (5′ primer P1 (TGCGAACTGAGATACGCAGGGA) and 3′ primer P3 (CATACTAGCCATTTTATGTG)). For this, 1µg of total gDNA was used to amplify the wild-type and integration loci in a T100 Thermal Cycler (Bio-Rad), with a PCR program consisting of 1 cycle at 95° C for 5 minutes, followed by 40 cycles of 95° C for 30 seconds, 50° C for 30 seconds and 68° C for 90 seconds, and a final cycle of 68° C for 10 minutes. The resulting PCR products were resolved in a 1% agarose gel and visualized in an Amersham® Imager 680. Clones were considered ablated for the *Plp1* gene by the presence of the amplification product corresponding to the correct integration of the modified target locus and absence of the wild-type locus.

### Immunizations and infections

Spz were obtained by the dissection of salivary glands of infected *Anopheles stephensi* mosquitoes from day 20 post blood meal onwards. All administrations of infectious or attenuated spz were done in 200µL of DMEM (Gibco) via retro-orbital (r.o.) route in mice lightly anesthetized with isoflurane (Isoflurane 1000mg/mL; Iso-Vet). RAS-based immunizations consisted of three administrations, on a weekly basis, of 10^4^ *Pb*WT, 15x10^4^ *PbSpect1*^-^, 15x10^4^ *PbPlp1*^-^ or 10^4^ *PbP36p*^-^ γ-irradiated spz (16,000 rads; Caesium137 γ-source, GammaCell 3000 Elan, Thetratronics). Non-immunized controls received equivalent amounts of salivary glands extract from uninfected *Anopheles stephensi* mosquitoes using the same volume and administration route. CPS-based immunizations consisted in three administrations, on a weekly basis, of 10^4^ *Pb*WT or 15x10^4^ *PbSpect1*^-^ infectious spz. Mice were administered azithromycin (AZ) (16mg/mL; Farmoz) in 200µL of saline (NaCl-0.9%; Braun), via intraperitoneal (i.p.) injection, on days 0, 1 and 2 post each immunization (CPS-AZ), or were given continuous access to water supplemented with chloroquine diphosphate salt (CQ) (0.288 mg/mL; Sigma) and glucose (15g/L; NZYtech), from days 1 to 6 post each immunization (CPS-CQ). Non-immunized controls received equivalent amounts of salivary glands extract from uninfected *Anopheles stephensi* mosquitoes using the same volume and administration route, and were exposed to AZ or CQ in doses equivalent to those of immunized cohorts. Mice from all immunization regimens and respective controls were challenged with 3x10^4^ *Pb*WT infectious spz, 28 days post priming.

### *In vivo* bioluminescence

Liver parasite burden at 48 h.p.c. was assessed by *in vivo* functional luciferase activity using the IVIS Lumina imaging system (PerkinElmer) and Living Image 3.0 software. Briefly, mice were administered D-Luciferin (180mg per Kg of body weight; PerkinElmer), diluted in 300μL of PBS, five minutes prior to being deeply anesthetized with a ketamine (100mg per Kg of body weight, Ketamidor; Richter Pharma) and xylazine (10mg per Kg of body weight, Rompum; Bayer) mixture via intraperitoneal (i.p.) injection. After 5 minutes, images were acquired with a 12.5cm field of view, medium binning factor and exposure time of 3 minutes. Quantitative analysis of bioluminescence in all experimental cohorts was performed by measuring the luminescence signal radiance using the Living Image 3.0 software.

The day of patency of blood stage infection was determined by detecting the presence of bioluminescence activity of circulating intraerythrocytic parasites using the Firefly Luciferase Assay kit 2.0 (Biotum) in a previously established blood bioluminescence assay^49^. Briefly, 5µL of blood was collected from days 3 to 12 post challenge from the tail vein of each mouse into 45µL of Lysis buffer 1x, and kept at -20° C until analysis. Bioluminescence was detected by adding 50µL of D-luciferin, previously diluted in Firefly Luciferase Assay Buffer (0.2mg/mL; Biotum), to 15µl of blood sample lysate and immediately measuring emitted luminescence, thrice, with an integration time of 100ms, using a Spark® multimode microplate reader (Tecan). Blood stage infection was considered patent when luminescence values increased above 100 for two consecutive days.

### ECM assessment

Mice survival and ECM establishment was evaluated twice daily, from day 6 post challenge onwards, by the emergence of neurological symptoms such as ataxia and paralysis, assessed by the ability of mice to self-right.

### Spz intrahepatic development and cell wounding assessment by immunofluorescence

*In vitro* infections were performed in HepG2 cells seeded on coverslips (12mm; Menzel-Gläser) at a density of 100,000 cells per well of a 24-well plate (Costar), cultured in DMEM medium supplemented with 10% Fetal Bovine Serum, 2 mM glutamine, and 100 U/mL penicillin-streptomycin (DMEMC) (all Gibco), and maintained at 37°C, 5% CO2, 24 hours prior to infection. For the assessment of cell wounding, rhodamine-labeled Dextran (0,5mg/mL; 10,000 Da, Lysine fixable; Molecular Probes) was added to each well immediately prior to infection with 1:1 ratio of cells to *Pb*WT or *PbPlp1*^-^ spz. Plates were incubated at 37°C, 5% CO2, and infection was halted 2 h.p.i.. For assessment of *in vitro* intrahepatic development, seeded HepG2 cells were infected with *Pb*WT or *Pbplp1*^-^ spz as before and infection was halted 48 h.p.i.. All cells were fixed with 4% paraformaldehyde (Santa Cruz Biotechnology Inc.).

*In vivo* infections for assessment of *Plasmodium*’s intrahepatic development were carried out as previously described. Forty-two hours post infection, mice were euthanized by CO_2_ narcosis, and livers were collected and fixed in 4% PFA overnight at 4 ° C. Livers were equilibrated in a 30% sucrose solution for 48 hours before embedding in optimal cutting temperature compound (Sakura Finetek), and frozen at -80°C. Tissue sections were serially cut in 10µm thick slices in a CM2050S cryostat (Leica) and mounted in glass slides (Marienfeld).

Fixed HepG2 cells were permeabilized with 0.1% (v/v) Triton X-100 (Sigma) before incubation with a blocking solution of 1% (weight/volume) of Albumin Bovine Fraction V (BSA) (NZYtech). Thawed slides with liver sections were permeabilized and blocked with a 5% BSA (w/v) (NZYtech) 0.3% (v/v) Triton X-100 (Sigma) in PBS.

For cell wounding assays, cells were incubated with Alexa Fluor 488-conjugated rabbit polyclonal αGFP antibody (4μg/mL; Invitrogen) and Hoechst 33342 (0.01μg/mL; Invitrogen), all diluted in blocking solution, for 90 minutes at room temperature. For *in vitro* and *in vivo* assessment of *Plasmodium*’s intrahepatic development, samples were incubated overnight at 4°C with goat αUIS4 polyclonal antibody (10μg/mL; Sicgen) followed by incubation with Alexa Fluor 568-conjugated donkey αgoat IgG (4μg/mL; Invitrogen), Alexa Fluor 488-conjugated rabbit αGFP polyclonal antibody (4μg/mL; Invitrogen) and Hoechst 33342 (0.01μg/mL; Invitrogen). For *in vivo* immunofluorescence staining, all antibodies were diluted in 5% BSA (w/v) (NZYtech) 0.3% (v/v) Triton X-100 (Sigma) in PBS. For all immunofluorescence assays, samples were mounted in Fluoromount G (Invitrogen) on microscope slides (Marienfeld). Mounted slides were allowed to dry in the dark at 4°C for, at least, 24 hours before imaging. Images were acquired using a LSM 880 with Airyscan microscope (Zeiss) at 20x (*in vivo* intrahepatic development), or at 63x (*in vitro* assays) magnification. EEF size was assessed using a macro in Fiji Image J (NIH).

### Isolation of hepatocytes for i*n vivo* cell wounding assessment

C57BL/6 mice received 160μg of rhodamine-labeled Dextran (10,000 Da, Lysine fixable; Molecular Probes) diluted in PBS immediately before inoculation of 5x10^5^ *Pb*WT or 5x10^5^ *PbP36p*^-^infectious spz, both via r.o. route. Two hours after, hepatocytes were isolated by a modified two-step collagenase perfusion method. ^50^ Briefly, livers were perfused *in situ* through vena cava, cutting the portal vein to allow for outflow, with 15mL of perfusion buffer (HBSS no Ca2^+^, no Mg2^+^, no Phenol Red (Gibco), supplemented with EDTA (0.5M; Gibco), and HEPES (25mM; Gibco)) pre-warmed at 42°C. Following, livers were perfused with 25mL of digestion buffer (HBSS with Ca2^+^, Mg2^+^, and Phenol Red (Gibco), supplemented with HEPES (25mM; Gibco) and Liberase Blendzyme 3 recombinant collagenase (0,04mg/mL; Roche Diagnostics)) pre-warmed at 42°C. After perfusion, liver tissue was gently dispersed in a single cell suspension using a pair of forceps. Single cell suspensions were passed through a 100μm filter, and liver parenchyma cells isolated in a 50% Percoll (Cytiva) density gradient, centrifuged at 70g for 10 minutes.

### Liver load assessment by quantitative reverse-transcriptase polymerase chain reaction (qRT-PCR)

C57BL/6 and BALB/c mice were euthanized and livers collected at the indicated time points following infection/challenge, homogenized in PureZOL RNA Isolation Reagent (Bio-Rad) using the Mini-Beadbeater-16 cell disrupter (BioSpect) for 3 minutes, and stored at -80°C until RNA extraction. Total RNA was extracted using the NZY Total RNA Isolation kit (NZYtech) according to the manufacturer’s instructions and quantified in a Nanodrop 1000 (ThermoScientific). *Pb18s* and *M. musculus Hprt* levels were assessed using TaqMan MGB probes (*Pb18s*: 5′ primer: AGGGAGCCTGAGAAATAG; 3′ primer: GTCACTACCTCTCTTATTTAGAA; probe: 6-FAM-ACCACATCTAAGGAAGGCAGCA-BHQ1; *mHprt*: 5′ primer: GAACCAGGTTATGACCTA; 3′ primer: TCTCCTTCATGACATCTC; probe: 6-FAM-TTCAGTCCTGTCCATAATCAGTCCAT-BHQ1). One-step NZYSpeedy RT-qPCR Probe kit (NZYtech) was used for tandem complementary DNA (cDNA) synthesis and amplification from 100ng of total RNA, in a RT-PCR 7500 Fast cycler (Applied Biosystems). The program consisted of 1 cycle at 50°C for 20 minutes, 1 cycle at 95°C for 5 minutes followed by 40 cycles of 95°C for 5 seconds and 60°C for 1 minute. The parameter threshold cycle (Ct) was used to quantify the relative expression of both parasite and host genes with the comparative ΔΔCt method, using *M. musculus Hprt* as the normalizing reference gene.

### *In vivo* αCD8 antibody administration

RAS-immunized C57BL/6 and BALB/c mice received 200mg of depleting αCD8 mouse monoclonal antibody (2.43; BioXCell) or IgG2b (LTF-2; BioXCell) diluted in saline via i.p. route, 24 days after priming, and 4 days prior to the administration of 3x10^4^ infectious *Pb*WT spz.

### Isolation and adoptive transfer of *Klrg1^Cre^Rosa26^tdTomato^*CD8 T cells

CD8 T cells from C57BL/6 *Klrg1^Cre^Rosa26^tdTomato^* mice were purified from a single cell suspension of spleen and lymph nodes, by labeling with biotin-conjugated αCD8 antibody (53-6.7; Biolegend) and positive selection with streptavidin MACS microbeads (Miltenyi Biotec), according to the manufacturer’s instructions. Recipient C57BL/6 mice received 5x10^6^ purified CD8 *Klrg1^Cre^Rosa26^tdTomato^* T cells, via r.o. route, in 200μL of PBS, 24 hours prior to priming.

### Isolation of splenocytes and tissue-infiltrating leukocytes

C57BL/6 and BALB/c mice were euthanized at indicated time points, transcardially perfused with 15mL of PBS, and livers and spleens were collected and homogenized in PBS. Liver-infiltrating leukocytes were isolated in a 35% Percoll (Cytiva) density gradient, centrifuged at 1360g for 20 minutes at room temperature. Following, erythrocytes were lysed and the total number of splenocytes and liver-infiltrating leukocytes was assessed in a 5-laser Cytek Aurora (Cytek Biosciences). *Ex-vivo* cytokine production of transferred *Klrg1^Cre^Rosa26^tdTomato^*CD8 T cells was assessed following culture with phorbol 12-myristate 13-acetate (PMA) (50ng/mL; Sigma) and ionomycin (500ng/mL; Sigma) for 4 hours at 37°C, 5% CO_2_, in the presence of brefeldin A (10μg/mL; Sigma).

### Flow cytometry

For high-parameter spectral flow cytometry analysis, liver-infiltrating leukocytes and splenocytes isolated from immunized cohorts were labeled with monoclonal antibodies specific for CD45 (30-F11; Biolegend), CD3 (17A2; Biolegend), NK1.1 (PK136; Biolegend), TCRɣδ (GL-3,GL3; eBiosciences), CD11b (M1/70; Biolegend), CD4 (RM4-5; BD Biosciences), CD8 (53-6.7; BD Biosciences), CD62L (MEL-14; Biolegend), CD69 (H1.2F3; Biolegend), CD44 (IM7; Biolegend), KLRG1 (2F1; eBiosciences), CD11a (M17/4; BD Biosciences), CD11c (N418; Cytek Biosciences), MHC II (M5/114.15.2; BD Biosciences), F4/80 (BM8; Biolegend) and Ly6C (HK1.4; Cytek Biosciences). Prior to surface staining, Fc receptors were blocked using a monoclonal antibody specific for CD16/CD32 (93; eBioscience). All antibodies were diluted in staining buffer (PBS with 2% FBS) and staining performed at 4°C. Dead cells were excluded using a live/dead fixable dye (Cytek Biosciences), diluted in PBS and performed at 4°C prior to incubation with surface or intracellular antibodies. Intracellular staining to evaluate cytokine production was performed using monoclonal antibodies specific for IFN-γ (XMG1.2; Biolegend) and Granzyme B (REA226; Miltenyi Biotec) after cell fixation and permeabilization using the FoxP3/Transcription Factor Staining Kit (Invitrogen), according to the manufacturer’s instructions. Samples were acquired in a 5-laser Cytek Aurora (Cytek Biosciences), and unmixing and compensation corrections were performed using single-color cell controls in SpectroFlo software V3.3.0 (Cytek Biosciences). Data analysis was performed inFlowJo V10.10.0 software (Tree Star Inc.).

For immunophenotyping of transferred *Klrg1^Cre^Rosa26^tdTomato^*CD8 T cells, liver-infiltrating leukocytes and splenocytes were labeled with monoclonal antibodies specific for CD45.1 (A20; BD Biosciences), CD45.2 (104; Biolegend), CD3 (17A2; Biolegend), NK1.1 (PK136; Biolegend), CD4 (RM4-5; BD Biosciences), CD8 (53-6.7; BD Biosciences), CD62L (MEL-14; Biolegend), CD69 (H1.2F3; Biolegend), CD44 (IM7; Biolegend) and KLRG1 (2F1; eBiosciences). Blocking of Fc receptors and dead cell exclusion was done as before. All antibodies were diluted in staining buffer. Intracellular staining was performed using monoclonal antibodies specific for IFN-γ (XMG1.2; Biolegend) and Granzyme B (REA226; Miltenyi Biotec) after cell fixation and permeabilization using the FoxP3/Transcription Factor Staining Kit (Invitrogen), according to the manufacturer’s instructions. Samples were acquired in a 5-laser Cytek Aurora (Cytek Biosciences), and unmixing and compensation corrections were performed using single-color cell controls in SpectroFlo software V3.3.0 (Cytek Biosciences). Data analysis was performed in FlowJo V10.10.0 software (Tree Star Inc.).

Progression of blood stage infection was evaluated by the daily assessment of parasitemia, i.e., the frequency of infected erythrocytes. For this, blood samples were collected from the tail vein in PBS and incubated for 10 minutes at 37°C with monoclonal antibodies specific for Ter-119 (TER-119; eBioscience) and CD45 (30-F11; Biolegend), and Hoechst 33342 (Invitrogen), all diluted in PBS. Samples were acquired on a BD LSRFortessa (BD Biosciences), and compensation was performed using single-color cell controls in FACSDiva software V6.2.0 (BD Biosciences). Data analysis was performed in FlowJo V10.10.0 software (Tree Star Inc.). Frequency of infected erythrocytes was determined by the percentage of Ter-119^+^ CD45^-^ Hoechst^+^ single cells.

To evaluate CD8 T cell depletion, circulating leukocytes, liver-infiltrating leukocytes and splenocytes were isolated at the indicated time points, and stained with monoclonal antibodies specific for CD45 (30-F11; Biolegend), CD3 (17A2; Biolegend), CD4 (GK1.5; Biolegend) and CD8 (53-6.7; Biolegend). Prior, Fc receptors were blocked using a monoclonal antibody specific for CD16/CD32 (93; eBioscience). All antibodies were diluted in PBS. Samples were acquired on a BD LSRFortessa (BD Biosciences), and compensation was performed using single-color cell controls in FACSDiva software V6.2.0 (BD Biosciences). Data analysis was performed in FlowJo V10.10.0 software (Tree Star Inc.).

To evaluate *in vivo* cell wounding, liver parenchyma cells were stained with a monoclonal antibody specific for CD31 (MEC13.3; Biolegend) and Live/Dead Fixable Violet Dead Cell (Invitrogen). Cells were acquired in a BD BD LSRFortessa(BD Biosciences), and compensation was performed using single-color cell controls in FACSDiva software V6.2.0 (BD Biosciences). Data analysis was performed in FlowJo V10.10.0.0 software (Tree Star Inc.).

### Hierarchical unsupervised analysis of spectral flow cytometry data

Analysis was performed using FlowJo V10.10.0 software (Tree Star Inc.). Abnormal events were excluded from all samples using the Peak Extraction and Cleaning Oriented Quality Control (PeacoQC) plugin. For the analysis of lymphocyte populations, a total of 80.000 live CD45^+^ CD3^+^ cells were randomly selected per condition, with equal numbers sampled from each mouse to ensure balanced representation. All cells (240,000 total) were concatenated into a single file. Dimensional reduction was performed using the UMAP plugin, followed by unsupervised FlowSOM hierarchical clustering based on the expression of CD8, CD4, NK1.1, TCRδ, CD62L, CD44, CD69, KLRG1 and CD11a. For the analysis of myeloid and natural killer cell populations, a total of 60.000 live CD45^+^ CD3⁻ cells expressing CD11b and/or NK1.1 were randomly selected from each condition, with equal numbers sampled from each mouse to ensure balanced representation. All cells (180,000 total) were concatenated into a single file. Dimensional reduction was again performed using the UMAP plugin. Hallmark markers used for hierarchical clustering with FlowSOM were MHCII, NK1.1, CD11b, F4/80, Ly6C and CD11c. For both concatenated files several numbers of clusters were tested. To avoid over-clustering, phenotypically similar clusters were manually merged. Ultimately, 14 and 7 were selected, respectively, as best representing inter-cluster differences in marker expression.

## ELISA

Titers of circulating αspz IgG antibodies were assessed in serum at the indicated time points, as previously described^41^. Briefly, serum was obtained by collecting 10µL of blood from the tail vein of mice into 90µL of PBS supplemented with heparin (10U/mL; Braun), followed by centrifugation for 2 minutes at 800g, at room temperature. Supernatants were stored at -20°C until analysis. Dissected *Pb*WT spz were purified by nycodenz (Alere Technologies) gradient, spz protein extracts were obtained by incubating 5x10^5^ purified spz in extraction buffer (150 mM NaCl, 20 mM Tris-HCl, 1% triton, 1 mM EDTA, pH= 7.5), and used to coat each 96-well Nunc Maxisorp ELISA plate (Sigma) overnight. Coated wells were incubated with blocking buffer (5% nonfat dry milk in PBS supplemented with 0,05% Tween20 (Sigma-Aldrich) (PBST)) for 15 minutes on ice, after which 100µL of diluted sera (1:1000) were added to each well and incubated for 2 hours at room temperature with continuous shaking. After washing with PBST, samples were incubated with HRP-conjugated goat αmouse IgG antibody (0.4μg/mL; Enzo Life Sciences) for 1 hour at room temperature with continuous shaking. Plates were washed with PBST and incubated with TMB Substrate Reagent Set (BD OptEIA, BD Biosciences), according to the manufacturer’s instructions. The colorimetric reaction was stopped after 10 minutes by adding 50µL of 2M H_2_SO_4_ and absorbance was measured at an optical density of 450nm using 570nm as a reference wavelength, in a Tecan Spark plate reader (Tecan).

### Statistical analysis

Statistically significant differences between two different groups were determined using the Log-rank (Mantel-Cox) test and the non-parametric two-tailed Mann-Whitney t-test, as indicated. All tests were carried out in GraphPad Prism software (V8.4.3.; GraphPad).

## Notes

### Competing Interest Statement

The authors have declared no competing interest.

## References

1. World Health Organization. World Malaria Report 2024. (2024).

2. Langhorne, J., Ndungu, F. M., Sponaas, A. M. & Marsh, K. Immunity to malaria: More questions than answers. Nat Immunol 9, 725–732 (2008).

3. Duffy, P. E. & Patrick Gorres, J. Malaria vaccines since 2000: progress, priorities, products. NPJ Vaccines 5, (2020).

4. Mota, M. M. et al. Migration of Plasmodium Sporozoites Through Cells Before Infection. Science (1979) 20, 141–144 (2001).

5. Risco-Castillo, V. et al. Malaria sporozoites traverse host cells within transient vacuoles. Cell Host Microbe 18, 593–603 (2015).

6. Datoo, M. S. et al. Efficacy of a low-dose candidate malaria vaccine, R21 in adjuvant Matrix-M, with seasonal administration to children in Burkina Faso: a randomised controlled trial. The Lancet 397, 1809–1818 (2021).

7. Datoo, M. S. et al. Safety and efficacy of malaria vaccine candidate R21/Matrix-M in African children: a multicentre, double-blind, randomised, phase 3 trial. The Lancet 403, 533–544 (2024).

8. The RTS, S. C. T. P. A Phase 3 Trial of RTS,S/AS01 Malaria Vaccine in African Infants. New England Journal of Medicine 367, 2284–2295 (2012).

9. The RTS, S. C. T. P. Efficacy and safety of RTS,S/AS01 malaria vaccine with or without a booster dose in infants and children in Africa: final results of a phase 3, individually randomised, controlled trial. The Lancet 386, 31–45 (2015).

10. Sissoko, M. S. et al. Safety and efficacy of a three-dose regimen of Plasmodium falciparum sporozoite vaccine in adults during an intense malaria transmission season in Mali: a randomised, controlled phase 1 trial. Lancet Infect Dis 22, 377–389 (2022).

11. Mwakingwe-Omari, A. et al. Two chemoattenuated PfSPZ malaria vaccines induce sterile hepatic immunity. Nature 595, 289–294 (2021).

12. Moita, D. et al. The effect of dosage on the protective efficacy of whole-sporozoite formulations for immunization against malaria. NPJ Vaccines 8, 182 (2023).

13. Kreutzfeld, O., Müller, K. & Matuschewski, K. Engineering of Genetically Arrested Parasites (GAPs) For a Precision Malaria Vaccine. Front Cell Infect Microbiol 7, (2017).

14. Goswami, D., Minkah, N. K. & Kappe, S. H. I. Designer Parasites: Genetically Engineered *Plasmodium* as Vaccines To Prevent Malaria Infection. The Journal of Immunology 202, 20–28 (2019).

15. Mo, A. X., McGugan, G. & Pesce, J. T. Meeting report: Expert consultation on late arresting replication competent (LARC) malaria sporozoite vaccine research & development. Vaccine 54, 127009 (2025).

16. Seder, R. A. et al. Protection Against Malaria by Intravenous Immunization with a Nonreplicating Sporozoite Vaccine. Science (1979) 341, 1359–1365 (2013).

17. Roestenberg, M. et al. A double-blind, placebo-controlled phase 1/2a trial of the genetically attenuated malaria vaccine PfSPZ-GA1. Sci Transl Med 12, (2020).

18. Ishino, T., Yano, K., Chinzei, Y. & Yuda, M. Cell-passage activity is required for the malarial parasite to cross the liver sinusoidal cell layer. PLoS Biol 2, (2004).

19. Ishino, T., Chinzei, Y. & Yuda, M. A Plasmodium sporozoite protein with a membrane attack complex domain is required for breaching the liver sinusoidal cell layer prior to hepatocyte infection. Cell Microbiol 7, 199–208 (2005).

20. Nganou-Makamdop, K., van Gemert, G.-J., Arens, T., Hermsen, C. C. & Sauerwein, R. W. Long Term Protection after Immunization with P. berghei Sporozoites Correlates with Sustained IFNγ Responses of Hepatic CD8+ Memory T Cells. PLoS One 7, e36508 (2012).

21. Butler, N. S., Vaughan, A. M., Harty, J. T. & Kappe, S. H. I. Whole parasite vaccination approaches for prevention of malaria infection. Trends Immunol 33, 247–254 (2012).

22. Matuschewski, K. Vaccines against malaria—still a long way to go. FEBS J 284, 2560–2568 (2017).

23. Friesen, J. & Matuschewski, K. Comparative efficacy of pre-erythrocytic whole organism vaccine strategies against the malaria parasite. Vaccine 29, 7002–7008 (2011).

24. Moita, D. & Prudêncio, M. Whole-sporozoite malaria vaccines: where we are, where we are going. EMBO Mol Med 16, 2279–2289 (2024).

25. Ishino, T., Chinzei, Y. & Yuda, M. Two proteins with 6-cys motifs are required for malarial parasites to commit to infection of the hepatocyte. Mol Microbiol 58, 1264–1275 (2005).

26. Abuga, K. M., Jones-Warner, W. & Hafalla, J. C. R. Immune responses to malaria pre-erythrocytic stages: Implications for vaccine development. Parasite Immunol 43, (2021).

27. Aguirre-Botero, M. C. et al. Late killing of Plasmodium berghei sporozoites in the liver by an anti-circumsporozoite protein antibody. Elife 14, (2025).

28. Keitany, G. J. et al. Immunization of Mice with Live-Attenuated Late Liver Stage-Arresting Plasmodium yoelii Parasites Generates Protective Antibody Responses to Preerythrocytic Stages of Malaria. Infect Immun 82, 5143–5153 (2014).

29. Guyach, S. E., Bryan, M. A. & Norris, K. A. Differences in antibody and immune responses between Balb/c and C57Bl/6 mice infected with *Trypanosoma cruzi* (129.5). The Journal of Immunology 182, 129.5–129.5 (2009).

30. Zeng, M., Nourishirazi, E., Guinet, E. & Nouri-Shirazi, M. The genetic background influences the cellular and humoral immune responses to vaccines. Clin Exp Immunol 186, 190–204 (2016).

31. Chu, Y. et al. CpG adjuvant enhances humoral and cellular immunity against OVA in different degrees in BALB/c, C57BL/6J, and C57BL/6N mice. Int Immunopharmacol 138, 112593 (2024).

32. Fernandez-Ruiz, D. et al. Liver-Resident Memory CD8+ T Cells Form a Front-Line Defense against Malaria Liver-Stage Infection. Immunity 45, 889–902 (2016).

33. Holz, L. E. et al. CD8+ T Cell Activation Leads to Constitutive Formation of Liver Tissue-Resident Memory T Cells that Seed a Large and Flexible Niche in the Liver. Cell Rep 25, 68–79.e4 (2018).

34. Doolan, D. L. & Hoffman, S. L. IL-12 and NK cells are required for antigen-specific adaptive immunity against malaria initiated by CD8+ T cells in the Plasmodium yoelii model. J Immunol 163, 884–92 (1999).

35. Miller, J. L., Sack, B. K., Baldwin, M., Vaughan, A. M. & Kappe, S. H. I. Interferon-Mediated Innate Immune Responses against Malaria Parasite Liver Stages. Cell Rep 7, 436–447 (2014).

36. Herndler-Brandstetter, D. et al. KLRG1+ Effector CD8+ T Cells Lose KLRG1, Differentiate into All Memory T Cell Lineages, and Convey Enhanced Protective Immunity. Immunity 48, 716–729.e8 (2018).

37. Renkema, K. R. et al. KLRG1+ Memory CD8 T Cells Combine Properties of Short-Lived Effectors and Long-Lived Memory. The Journal of Immunology 205, 1059–1069 (2020).

38. Goswami, D., Minkah, N. K. & Kappe, S. H. I. Designer Parasites: Genetically Engineered *Plasmodium* as Vaccines To Prevent Malaria Infection. The Journal of Immunology 202, 20–28 (2019).

39. Vanderberg, J., Nussenzweig, R. & Most, H. Protective immunity produced by the injection of x-irradiated sporozoites of Plasmodium berghei. V. In vitro effects of immune serum on sporozoites. Mil Med 134, 1183–90 (1969).

40. Sano, G. et al. Swift Development of Protective Effector Functions in Naive Cd8+ T Cells against Malaria Liver Stages. J Exp Med 194, 173–180 (2001).

41. Nunes-Cabaço, H., Moita, D., Rôla, C., Mendes, A. M. & Prudêncio, M. Impact of Dietary Protein Restriction on the Immunogenicity and Efficacy of Whole-Sporozoite Malaria Vaccination. Front Immunol 13, (2022).

42. Behr, F. M., Chuwonpad, A., Stark, R. & van Gisbergen, K. P. J. M. Armed and Ready: Transcriptional Regulation of Tissue-Resident Memory CD8 T Cells. Front Immunol 9, (2018).

43. Pritzl, C. J., Daniels, M. A. & Teixeiro, E. Interplay of Inflammatory, Antigen and Tissue-Derived Signals in the Development of Resident CD8 Memory T Cells. Front Immunol 12, (2021).

44. Ma, M., Jiang, W. & Zhou, R. DAMPs and DAMP-sensing receptors in inflammation and diseases. Immunity 57, 752–771 (2024).

45. Häger, S. C. et al. Short-term transcriptomic response to plasma membrane injury. Sci Rep 11, 19141 (2021).

46. Kazmin, D. et al. Systems analysis of protective immune responses to RTS,S malaria vaccination in humans. Proceedings of the National Academy of Sciences 114, 2425–2430 (2017).

47. Janse, C. J. et al. High efficiency transfection of Plasmodium berghei facilitates novel selection procedures. Mol Biochem Parasitol 145, 60–70 (2006).

48. Billker, O. et al. Calcium and a Calcium-Dependent Protein Kinase Regulate Gamete Formation and Mosquito Transmission in a Malaria Parasite. Cell 117, 503–514 (2004).

49. Zuzarte-Luis, V., Sales-Dias, J. & Mota, M. M. Simple, sensitive and quantitative bioluminescence assay for determination of malaria pre-patent period. Malar J 13, 15 (2014).

50. Afriat, A. et al. A spatiotemporally resolved single-cell atlas of the Plasmodium liver stage. Nature 611, 563–569 (2022).

